# Microtubule rescue at midzone edges promotes overlap stability and prevents spindle collapse during anaphase B

**DOI:** 10.1101/2021.08.06.455369

**Authors:** Manuel Lera-Ramirez, François J. Nédélec, Phong T. Tran

**Affiliations:** Institut Curie, PSL Research University, Sorbonne Université, CNRS, UMR 144, F-75005 Paris, France; Sainsbury Laboratory, Cambridge University, 47 Bateman Street, Cambridge, CB2 1LR, UK; University of Pennsylvania, Department of Cell and Developmental Biology, Philadelphia, PA 19104, USA

## Abstract

During anaphase B, molecular motors slide interpolar microtubules to elongate the mitotic spindle, contributing to the separation of chromosomes. However, sliding of antiparallel microtubules reduces their overlap, which may lead to spindle breakage, unless microtubules grow to compensate sliding. How sliding and growth are coordinated is still poorly understood. In this study, we have used the fission yeast *S. pombe* to measure microtubule dynamics during anaphase B. We report that the coordination of microtubule growth and sliding relies on promoting rescues at the midzone edges. This makes microtubules stable from pole to midzone, while their distal parts including the plus ends alternate between assembly and disassembly. Consequently, the midzone keeps a constant length throughout anaphase, enabling sustained sliding without the need for a precise regulation of microtubule growth speed. Additionally, we found that in *S. pombe*, which undergoes closed mitosis, microtubule growth speed decreases when the nuclear membrane wraps around the spindle midzone.

## Introduction

The mitotic spindle is a bipolar assembly of microtubules, motors and microtubule associated proteins (MAPs), that orchestrates chromosome segregation. During prophase and metaphase, kinetochore microtubules capture and biorient chromosomes, and in anaphase A they transport chromosomes from the cell equator to the spindle poles. In anaphase B, the spindle elongates to further separate the chromatids. In certain cells (e.g. PtK2 cells) [1], cortical pulling on astral microtubules is thought to drive spindle elongation during anaphase B. However, in most biological systems studied so far [2–6], this is mainly driven by molecular motors, which slide interpolar microtubules at the midzone, the central spindle region where interpolar microtubules coming from opposite poles are crosslinked antiparallely by members of the PRC1/Ase1 family [7–9] (Fig. 1A). Importantly, sliding shortens the overlap between microtubules, such that microtubules must continuously elongate to sustain sliding [6, 10]. Additionally, the midzone length remains roughly constant during anaphase B [3, 11], indicating that net polymerisation and sliding closely match *in vivo*. How this happens is still poorly understood. Pioneering studies found that purified algae spindles would slide if tubulin was available for microtubules to grow, but would stop sliding in the absence of soluble tubulin [12]. This showed that sliding can be limited by microtubule growth. In higher eukaryotes, kinesin-4 suppresses the dynamics of interpolar microtubules and could induce this growth-limited regime [11, 13, 14]. However, depletion of kinesin-4, which leads to highly dynamic interpolar microtubules [11], does not have an impact on anaphase spindle elongation velocity [4]. Yeasts lack kinesin-4, but their final spindle length is reduced if microtubule dynamics are suppressed by nocodazole treatment or deletion of the microtubule polymerase Stu2 [15], indicating that suppressing microtubule dynamics can arrest sliding. Comparable results have been obtained in HeLa cells when depleting TACC3 [16]. Therefore, growth-limited sliding can happen in a wide range of organisms when microtubule dynamics are suppressed, but it is not known whether it occurs in unperturbed spindles. An alternative mechanism for the coordination of sliding and growth has been recently proposed: a motor could set both the polymerisation and sliding speed of interpolar microtubules [17]. In fission yeast, this would be Klp9 (kinesin-6), which slides microtubules during anaphase [3] and promotes microtubule polymerisation in monopolar spindles [17]. Klp9 and the related kinesin-5 also regulate microtubule dynamics *in vitro* [17, 18].

**Fig. 1:**
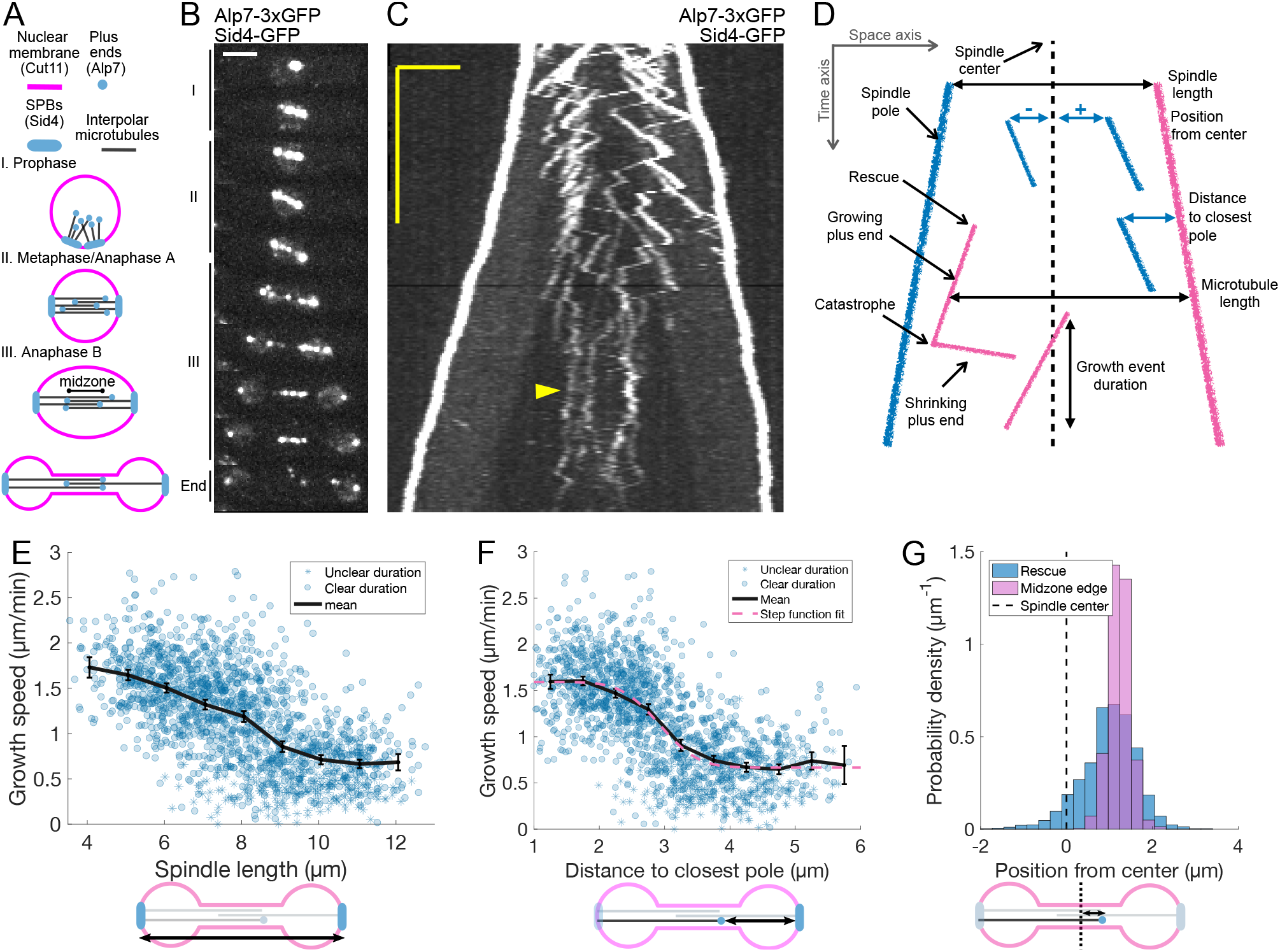
Characterization of microtubule dynamics during *S. pombe* anaphase. (**A**) The three phases of mitosis in the *S. pombe* spindle. Prophase (I), metaphase/anaphase A (II) and anaphase B (III). Names of proteins in parenthesis indicate the markers used to label the different components. (**B**) Time-lapse images of a mitotic spindle in a cell expressing Alp7-3xGFP and Sid4-GFP. Mitotic phases are indicated on the left. Time between images is 3 minutes, scale bar 3 μm. (**C**) Kymograph of a mitotic spindle during anaphase in a cell expressing Alp7-3xGFP and Sid4-GFP. Time is in the vertical axis (scalebar 5 minutes), and space is in the horizontal axis (scalebar 2 μm). Arrowhead marks a microtubule growth event in which the start and finish cannot be unambiguously determined. (**D**) Elements that can be identified in a kymograph (left) and the derived measurements (right). Pink plus ends have their minus end at the pink pole, and blue plus ends have their minus end at the blue pole. (**E-F**) Microtubule growth speed as a function of spindle length (E) or distance from the plus-end to closest pole (F) at rescue (or first point if rescue could not be exactly determined). Microtubule growth events of clear duration are shown as round dots, others as stars. Thick black lines represent average of binned data, error bars represent 95% confidence interval of the mean. Pink line in (F) represents a fit of the data to an error function. (**G**) Histogram showing the distribution of the position of rescues with respect to the spindle center in cells expressing Sid4-GFP and Alp7-3xGFP (blue), and position of midzone edges with respect to the spindle center in cells expressing mCherry-Atb2 and Cls1-3xGFP (pink, see Fig. 1-Supplement 1E, F). Dashed lines represent spindle center. Cartoons below the axis in (E-G) illustrate how the magnitudes represented are measured. Data shown in blue in E-G comes from 1671 growth events (119 cells), from 13 independent experiments (same data as wild type in Fig. 2G, J, K combined). Data shown in pink in G comes from 832 midzone length measurements during anaphase, from 60 cells in 10 independent experiments.

The contributions of microtubule nucleation and rescue to coordination of sliding and growth are less understood. Microtubule nucleation could enable sustained sliding by continuously replenishing microtubules in the overlap, similarly to what happens in *Xenopus* metaphase [19]. However, chromosome segregation during anaphase can occur in the absence of nucleation [20–22]. On the other hand, microtubule rescue is required during anaphase in *Xenopus* and *S. pombe*, and in the absence of the rescue promoting factor CLASP, spindle microtubules fully depolymerise at anaphase onset [23, 24]. In various species, CLASP orthologues are recruited to the spindle midzone by direct interaction with PRC1/Ase1 [24–26], which may enrich rescues at the midzone.

Finally, yeasts undergo closed mitosis: their nuclear membrane does not disassemble during mitosis, and in anaphase B, it constricts to form a dumbbell shape with a thin nuclear bridge around the spindle [27] (Fig. 1A). Recent studies have highlighted the interplay between the nuclear membrane and the central spindle [27–29]. For example, Ase1 midzone crosslinkers are required for the sorting of nuclear pore complexes to the nuclear membrane bridge [29], and for the nuclear bridge to act as a diffusion barrier between daughter nuclei [30]. Such a cross-talk between the nuclear membrane and the central spindle might affect microtubule dynamics.

The key to understand how microtubule sliding and polymerisation are coordinated is to directly measure anaphase B microtubule dynamics *in vivo*. FRAP has been used to infer certain aspects in cells expressing fluorescent tubulin, which established that microtubule turnover is lower in anaphase B when compared to metaphase in *Drosophila* and yeast [31–34]. These experiments also showed that no microtubule nucleation occurs during anaphase B in yeast, and that interpolar microtubules are maintained through rescues [35]. In HeLa cells, the turnover of EB1, a protein that binds to the tips of growing microtubules, decreases with anaphase progression [36], suggesting that there might be changes of microtubule dynamics within anaphase B. Finally, using fluorescently labelled tubulin, microtubule dynamics were observed in fission yeast spindles, but only at very late anaphase, when the spindle is composed of approximately four microtubules [37]. To our knowledge, no publication to date provides direct measurements of interpolar microtubule dynamics throughout anaphase B in any organism.

In this study, we have measured microtubule dynamics during anaphase B in *S. pombe* cells. We found that mi-crotubule growth speed decreases when the nuclear membrane wraps around the spindle midzone. Our observations support a model in which coordination of microtubule growth and sliding is not based on a precise regulation of microtubule growth speed, but instead relies on promoting rescues at the midzone edges. This makes microtubules stable from pole to midzone, as only their distal parts including the plus ends alternate between assembly and disassembly. Consequently, the midzone persists throughout anaphase, enabling sustained sliding.

## Results

### Live-imaging of S. pombe cells expressing Alp7-3xGFP allows to visualise microtubule dynamics during anaphase B

To measure microtubule dynamics during anaphase, we constructed *S. pombe* strains expressing Alp7 tagged with 3xGFP at the C-terminus from its endogenous locus. Alp7 is the *S. pombe* orthologue of mammalian Transforming Acidic Coiled-Coil (TACC) [38], and forms a complex with the XMAP215 orthologue Alp14, a microtubule polymerase [39]. The Alp7/Alp14 complex localises to Spindle Pole Bodies (SPBs) and microtubule plus ends [38]. Additionally, our strains expressed the SPB marker Sid4-GFP (Fig. 1A, B). Live-imaging of these cells during anaphase B using Structured Illumination Microscopy (SIM) produced movies where microtubule plus ends could be resolved (Movie 1). From such movies, we constructed kymographs where the duration and velocity of microtubule growth and shrinkage events can be measured (Fig. 1C, D). Given that in *S. pombe* all the minus ends of interpolar microtubules are located at the SPBs [40, 41], microtubule length can be measured as the distance between the plus end and the pole located opposite to the direction of growth (Fig. 1D). Note that since microtubules slide as the SPBs move apart, vertical comets in kymographs (Fig. 1C) do not correspond to non-growing microtubules, but rather microtubules that grow at a speed matching the sliding speed. In the kymographs some comets superimpose, so we could not count the total number of microtubules and it was not always possible to determine exactly where a microtubule starts or stops growing, especially at late stages of anaphase B (Fig. 1C, arrowhead). Nevertheless, the positions of SPBs and multiple microtubule tips could be determined without ambiguity, and thus the growing speed of microtubules could be measured reliably.

### Microtubule growth velocity decreases with anaphase B progression

We then proceeded to measure the parameters of microtubule dynamics from kymographs. We used spindle length as a proxy for anaphase B progression, as spindle elongation is very stereotypical in *S. pombe* (Fig. 1-Supplement 1A). Three phases can be observed corresponding to prophase (I), metaphase/anaphase A (II) and anaphase B (III) (Fig. 1A, B, Fig. 1-Supplement 1A). We found that the duration of microtubule growth events did not change during anaphase B and was on average 52±23 seconds (Fig. 1-Supplement 1B, C), significantly shorter than in interphase, where growth events last more than 120 seconds on average [42, 43]. Microtubule shrinking speed did not change during anaphase either (Fig. 1-Supplement 1D), and was on average 3.56±1.75 μm/min, also lower than in interphase (~8 min/μm) [42, 43].

Surprisingly, microtubule growth velocity decreased during anaphase B (Fig. 1C, E). Furthermore, this decrease was not gradual, and we observed two populations of microtubules (fast and slow growing) characteristic of early and late anaphase. In some cells, all microtubules seemed to switch to the slow growing phase simultaneously (Fig. 1C), while in others fast and slow growing microtubules co-existed (Fig. 2A). A good way to visualise the two populations is to plot microtubule growth speed as a function of the distance between the plus end and the closest pole at the time of rescue (Fig. 1D, F). This distance increases for all microtubules as the spindle elongates, and is higher for microtubules that are rescued closer to the center for a given spindle length, hence providing a reference frame aligned with the spindle. On such a plot, the data points visibly cluster in two separate clouds and the variation of growth speeds can be fitted by an error function (Fig. 1F), which would be characteristic of a system in which a transition between two states occurs at a given distance to the pole. The microtubule growth velocities of these two states, extracted from the fit, were 1.60 μm/min and 0.67 μm/min, both below the average growth speed of interphase microtubules (~2.3 μm/min) [42, 43].

**Fig. 2:**
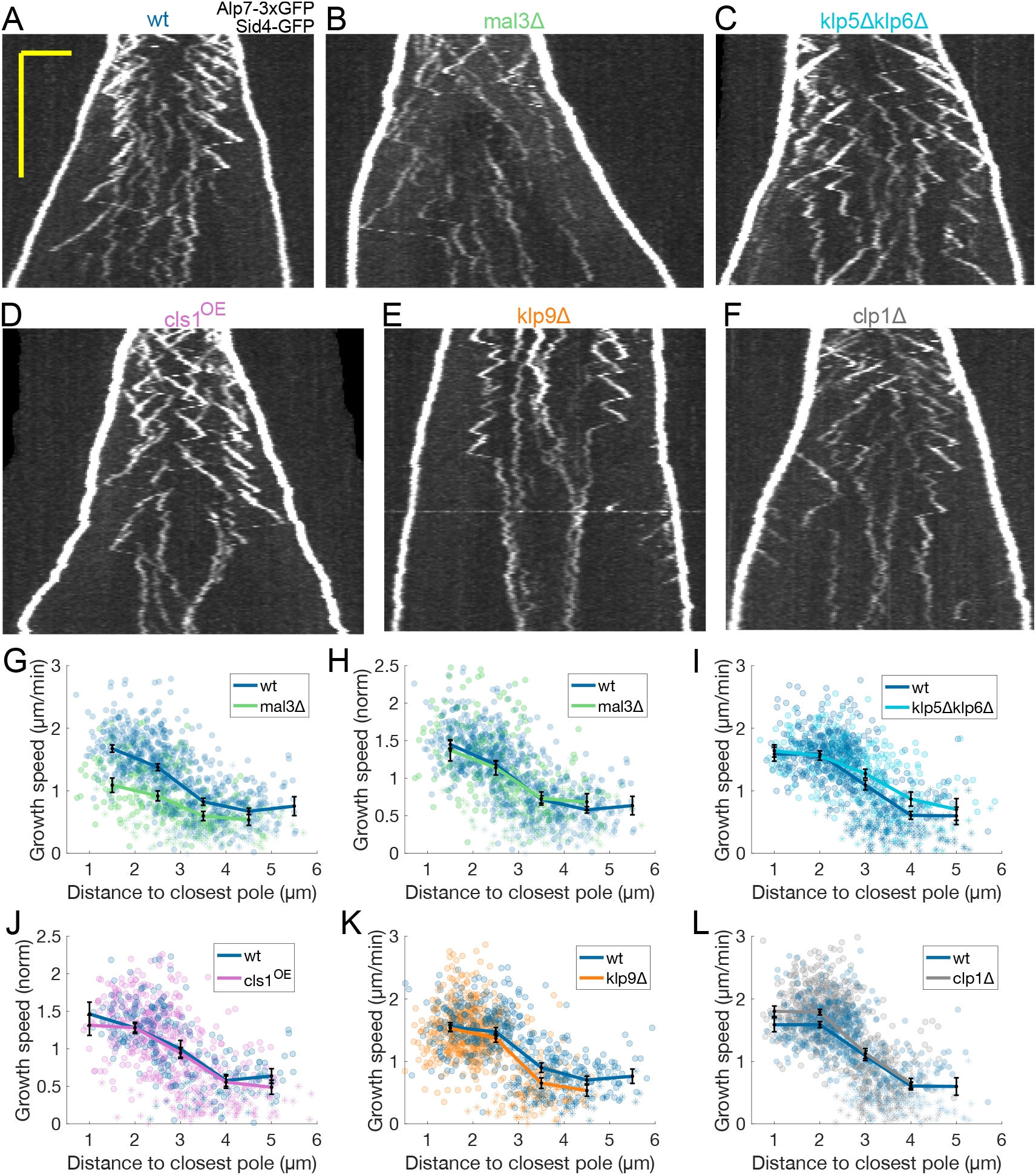
Transition from fast to slow microtubule growth occurs in the absence of known anaphase MAPs. (**A-F**) Kymographs of anaphase mitotic spindles in cells expressing Alp7-3xGFP and Sid4-GFP in different genetic backgrounds, indicated on top of each kymograph. Time is in the vertical axis (scalebar 5 minutes), and space is in the horizontal axis (scalebar 2 μm). (**G-L**) Microtubule growth speed as a function of the distance between the plus end and the closest pole at rescue (or first point if rescue could not be exactly determined) in the genetic backgrounds indicated by the legends, and shown in (A-F). Microtubule growth events of clear duration are shown as round dots, others as stars. Thick lines represent average of binned data, error bars represent 95% confidence interval of the mean. In (H) and (J), microtubule growth speed in each condition is normalised by the average value. Number of microtubule growth events shown: (G-H) 836 (59 cells) wt, 301 (37 cells) kymographs mal3Δ from 6 experiments (I) 589 (47 cells) wt, 405 (29 cells) klp5Δklp6Δ from 5 experiments (J) 248 (22 cells) wt, 532 (40 cells) from cls1^OE^ from 3 experiments (K) 531 (36 cells) wt, 610 (40 cells) klp9Δ from 4 experiments (L) 589 (47 cells) wt, 753 (58 cells) clp1Δ from 5 experiments.

### Microtubule rescues occur most often at midzone edges

In our dataset, most microtubule rescues occurred near the spindle midzone, where Cls1 (a MAP required for microtubule rescue, orthologue of CLASP) localises [24]. To describe the rescue distribution, we measured the positions of rescues with respect to the spindle center, considering the orientation of the microtubule to define negative and positive positions (Fig. 1D). This revealed that rescues did not occur uniformly along the midzone, since 93% of the rescues happened at positive positions (Fig. 1G). Moreover, the positions where most rescues happened coincided with the edge of the midzone, measured from Cls1-3xGFP signal in another strain (Fig. 1G and Fig. 1-Supplement 1E-F).

We observed similar microtubule dynamics in strains expressing GFP-mal3 (orthologue of EB1) [44], indicating that Alp7-3xGFP is a reliable marker for microtubule dynamics (Fig. 1-Supplement 2). Notably, GFP-Mal3 signal was lost from the spindle at late anaphase B (Fig. 1-Supplement 2A, B), making it impossible to track plus ends. This was not due to bleaching, as the GFP-Mal3 signal was not observed even if imaging was started at late anaphase B (data not shown). In contrast, Alp7 remains at microtubule plus ends at late anaphase. Moreover, Alp7 is a better marker for microtubule dynamics during anaphase B than the conventional EB1 marker, because it marks both growing and shrinking ends, and has a stronger signal.

In summary, our novel imaging approach shows that microtubule dynamics in anaphase are different from those previously reported for interphase. Additionally, at mid-anaphase microtubules transition from a fast growing state to a slow growing state. Finally, microtubule rescues happen with the highest probability at the midzone edge.

### Transition from fast to slow microtubule growth occurs in the absence of known anaphase MAPs

To investigate the intriguing transition from fast to slow microtubule growth velocity seen in anaphase B, we next examined various candidate MAPs associated with spindle microtubules. In *S. cerevisiae*, two MAPs regulate microtubule dynamics during anaphase: Bim1 (EB1) [45] and Kip3 (kinesin-8) [15].

Bim1 tracks the tips of growing microtubules and promotes microtubule growth [46]. At late anaphase, Bim1 is phosphorylated by Aurora B [45], which reduces its microtubule growth promoting activity [45]. In *S. pombe*, Mal3 (Bim1/EB1 orthologue) is also phosphorylated in a cellcycle dependent manner [47], and mutant Mal3 constructs mimicking this phosphorylation have lower microtubule growth promoting activity *in vitro* than wild-type Mal3 [47]. This, combined with our observation that GFP-Mal3 leaves the spindle at late anaphase B (Fig. 1-Supplement 2A) made it plausible that a combination of phosphorylation and unbinding of Mal3 could lead to the decrease in microtubule growth speed. To test this possibility, we used cells deleted for mal3. As expected, considering that Mal3 promotes microtubule growth [44], mal3Δ cells exhibited lower microtubule growth speed throughout anaphase B (Fig. 2B, G). However, microtubule growth velocities normalised by the mean in each condition were distributed similarly in wild-type and mal3Δ cells (Fig. 2H), suggesting that Mal3 is not required for the reduction of microtubule growth speed in anaphase B.

*S. cerevisiae* Kip3 (kinesin-8) is a plus end directed motor that localises to the anaphase B spindle and suppresses microtubule dynamics in a length-dependent manner [15, 48]. In *S. pombe* interphase, the kinesin-8 heterodimer Klp5/Klp6 accumulates at plus ends in a length-dependent manner, promoting catastrophe [43], so Klp5/Klp6 could trigger the transition from fast to slow microtubule growth during anaphase. We tested this hypothesis by deleting klp5 and klp6. As expected, klp5Δklp6Δ cells exhibited slightly longer microtubule growth events (Fig. 2-Supplement 1A). However, the distribution of microtubule growth speed as a function of distance from the plus end to the closest pole at rescue was not very different from wild-type cells (Fig. 2C, I), indicating that the decrease in microtubule growth speed is independent of Klp5/Klp6.

We next tested the human CLASP orthologue Cls1, a MAP that localises to the spindle midzone. Cls1 is required for microtubule rescues to occur during anaphase in *S. pombe* [24], and it decreases microtubule growth speed in a dose dependent manner [24, 49]. We measured Cls1 levels on the spindle and found that they remained constant throughout anaphase B (Fig. 2-Supplement 1B). However, as the number of microtubules in the spindle decreases with anaphase B progression [40, 41], the density of Cls1 on microtubules increases (Fig. 2-Supplement 1C), and this might reduce growth speed. Cls1 is an essential gene and cannot be deleted [50]. To study its potential role in regulating microtubule growth, we altered the levels of Cls1 by placing the gene under the control of a P81nmt1 promoter (cls1^off^), which reduced the amount of Cls1 on the spindle by approximately 60% (Fig. 2-Supplement 1B, C), or a P1nmt1 promoter (cls1^OE^), which led to a more than 4-fold increase in the amount of Cls1 on the spindle (Fig. 2-Supplement 1D, E). Consistent with its growth suppression activity [24, 49], reducing Cls1 levels slightly increased microtubule growth speed (Fig. 2-Supplement 1F), and overexpression of Cls1 slightly reduced growth speed (Fig. 2D, Fig. 2-Supplement 1G). However, these changes in microtubule growth speed were minor compared to the differences on Cls1 density (Fig. 2-Supplement 1C, E). Additionally, as in mal3Δ cells, the normalised growth speeds were distributed similarly to wild type in both cls1^off^ and cls1^OE^ cells (Fig. 2J, Fig. 2-Supplement 1H). In other words, the transition of microtubule growth speed occurred across a wide range of Cls1 levels, arguing against a role of Cls1 in this transition.

Finally, we tested the role of Klp9 (kinesin-6), the main driver of microtubule sliding during anaphase B [3]. Klp9 promotes microtubule growth *in vitro* and is required for the elongation of bundles of parallel microtubules present in monopolar spindles that undergo metaphase to anaphase transition [17]. Notably, the bundle elongation velocity matches the microtubule sliding speed in bipolar spindles, suggesting that Klp9 might set both microtubule growth and sliding speed in bipolar spindles. To test this hypothesis, we measured microtubule growth velocity during anaphase in cells deleted for klp9. As reported previously [3], klp9 deletion reduced spindle elongation velocity (Fig. 2-Supplement 1I). Interestingly, the decrease in microtubule growth velocity was delayed in klp9Δ cells with respect to wild-type (Fig. 2-Supplement 1J, K), while the distribution of microtubule growth speed as a function of the distance from the plus end to the closest pole at rescue was similar (Fig. 2E, K). This could indicate that the transition in microtubule growth velocity depends primarily on the position of the plus end of the microtubule when it is rescued. Alternatively, klp9 deletion could delay the activation of a regulatory network that decreases microtubule growth speed. In any case, Klp9 is not required for the transition in microtubule growth speed to occur.

Recruitment of Klp9 to the spindle midzone relies on the dephosphorylation of its Cdk1 phosphosites at anaphase onset [3]. This dephosphorylation occurs through two independent pathways: one involves the phosphatase Clp1 (orthologue of Cdc14) [3], which dephosphorylates multiple Cdk1 substrates during anaphase [51, 52], the other involves Dis1 [17], a member of the XMAP215 microtubule polymerase family [53]. Hence, microtubule growth speed could be regulated by Dis1 itself or by downstream effectors of Dis1 or Clp1. clp1Δ and dis1Δ cells had an identical phenotype to klp9Δ cells: their spindle elongation velocity was slower during anaphase, but showed no difference in the distribution of microtubule growth speed as a function of distance from the plus end to the closest pole at rescue (Fig. 2F, L, Fig. 2-Supplement 1L), indicating that neither the dephosphorylation of Cdk1 phosphosites mediated by Clp1 nor the activity of Dis1 are required for the transition from fast to slow microtubule growth velocity to occur during anaphase B.

In summary, our experiments show that the transition from fast to slow microtubule growth during anaphase occurs in the absence of multiple MAPs associated with the mitotic spindle known to affect microtubule dynamics in a different context or organism. Interestingly, the distribution of microtubule growth speed as a function of the distance from the plus end to the closest pole at rescue is maintained even in cells where spindle elongation is slower, suggesting that microtubule growth velocity during anaphase B could be affected by spatial cues associated with the spindle.

### Microtubules grow slower when they enter the nuclear membrane bridge formed at the dumbbell transition

Recent reports provide increasing evidence of a crosstalk between the spindle midzone and the nuclear membrane bridge that forms after the dumbbell transition in closed mitosis (Fig. 1A) [27–29]. Imaging strains expressing the nuclear membrane marker Cut11-mCherry (Fig. 3A), we noticed that the slow growing microtubules were the ones inside the nuclear membrane bridge (Fig. 3B, C). To verify this, we measured microtubule growth speed in wild-type cells, in cells overexpressing Klp9 (klp9^OE^), and in cells expressing the cell-cycle mutant allele cdc25-22, which all have different sizes and spindle elongation dynamics [54]. klp9^OE^ cells were longer than wild type cells, and their spindle elongation velocity was higher (Fig. 3-Supplement 1A-C), while cdc25-22 cells were bigger than wt and klp9^OE^ cells, and displayed an intermediate spindle elongation velocity (Fig. 3-Supplement 1A-C). Wild-type, klp9^OE^and cdc25-22 nuclei underwent dumbbell transition at different spindle lengths (Fig. 3-Supplement 1D), and their distribution of microtubule growth speed as a function of time, spindle length or distance from the plus end to the closest pole at rescue were different (Fig. 3-Supplement 1E-H). However, regression analysis showed that if we categorised microtubule growth events depending on whether they occurred ‘before’ the dumbbell transition, and ‘inside’ or ‘outside’ the nuclear membrane bridge (cartoons in Fig. 3D), microtubule growth velocity in each category was not different for wt, klp9^OE^ and cdc25-22 cells (Fig. 3D), and 48% of the variability was explained by this categorisation (Table S1). The 8% decrease in growth speed observed between ‘before’ and ‘outside’ events was minor compared to the 58% reduction when comparing ‘before’ and ‘inside’ (Fig. 3D). This categorisation clarifies our plots of microtubule growth speed as a function of the distance from the plus end to the closest pole at rescue (Fig. 1F), as this distance is equivalent to the distance from the plus end to the nuclear membrane bridge edge (Fig. 3E).

**Fig. 3:**
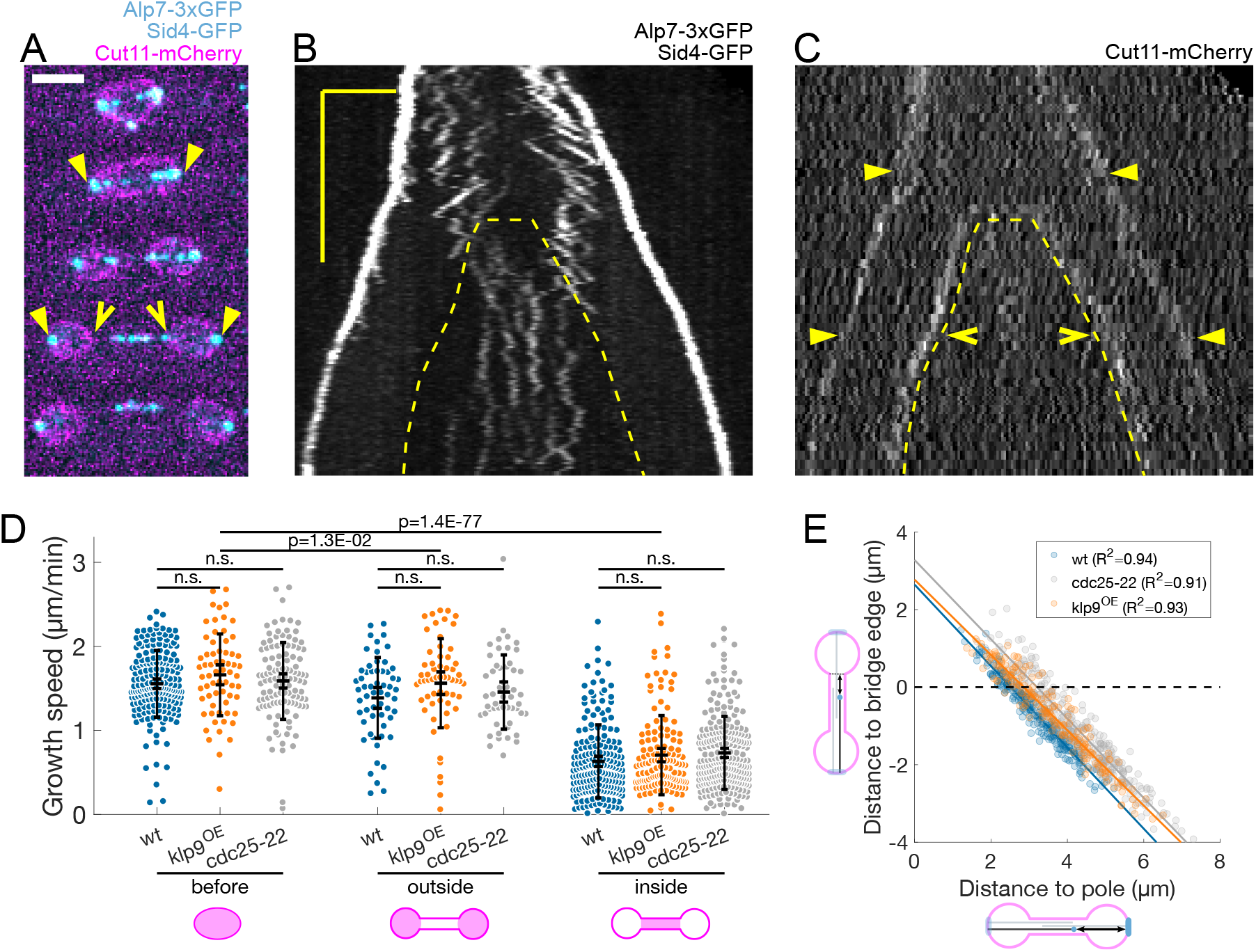
Microtubules grow slower when they enter the nuclear membrane bridge formed at the dumbbell transition. (**A**) Time-lapse images of an anaphase B mitotic spindle in a cell expressing Alp7-3xGFP, Sid4-GFP (cyan) and Cut11-mCherry (magenta). Time between images is 3 minutes, scale bar 3 μm. Filled arrowheads denote spindle poles, and empty arrowheads denote limits of the nuclear membrane bridge. Equivalent positions are marked in the kymograph in (C). (**B-C**) Kymographs of an anaphase B mitotic spindle in a cell expressing Alp7-3xGFP, Sid4-GFP (B) and Cut11-mCherry (C). Time is in the vertical axis (scalebar 5 minutes), and space is in the horizontal axis (scalebar 2 μm). Dashed lines outline the nuclear membrane bridge formed after the dumbbell transition (see Fig. 1A). See legend of (A) for arrowheads. (**D**) Microtubule growth speed in wild-type (blue), klp9^OE^ (orange) and cdc25-22 (grey) cells. Events are categorised according to whether rescue occurred before the dumbbell transition, and inside or outside the nuclear membrane bridge (see cartoons under x-axis). Error bars represent 95% confidence interval of the mean and standard deviation. p-values are calculated with regression analysis, ‘n.s’ (not significant) indicates p>0.05. See Table S1. (**E**) Distance from the plus-end to the nuclear membrane bridge edge at rescue as a function of distance from the plus-end to the closest pole at rescue. Dots represent individual microtubule growth events, with colour code as in (D). Lines represent first-degree polynomial fit to the data in each condition, of which the R^2^ is shown in the legend. Number of events: 442 (30 cells) wt, 260 (27 cells) klp9^OE^, 401 (35 cells) cdc25-22, from 3 independent experiments.

From this direct imaging we conclude that microtubules grow slower when they enter the nuclear membrane bridge formed by the dumbbell transition.

### Preventing the dumbbell transition abolishes the switch from fast to slow microtubule growth

To check whether the nuclear membrane bridge is indeed causing microtubule growth velocity to decrease at mid-anaphase, we prevented the nuclear membrane dumbbell transition in two different ways.

First, we inhibited Aurora B using an analogue sensitive allele (ark1-as3) [55], which often led to failed chromosome segregation [56], and spindles that elongated without undergoing dumbbell transition (Fig. 4B). To distinguish the direct effects of Aurora B inhibition from the effects of preventing the dumbbell transition, we added the analogue 1NM-PP1 and imaged cells immediately. Some of the cells were in anaphase B when the analogue was added, and displayed normal chromosome segregation (‘ark1-as3 normal’), others failed chromosome segregation and did not undergo dumbbell transition (‘ark1-as3 abnormal’). In wild-type and ‘ark1-as3 normal’ cells, microtubules grew slower inside the nuclear membrane bridge (Fig. 4I, Table S1), but in ‘ark1-as3 abnormal’ cells microtubule growth velocity was the same outside or inside the nuclear membrane tubes formed at the spindle poles (Fig. 4F, I, Fig. 4-Supplement 1B, Table S1). This data shows that the transition from fast to slow microtubule growth can happen upon Aurora B inactivation, but not if chromosome segregation fails and the dumbbell transition is prevented.

**Fig. 4:**
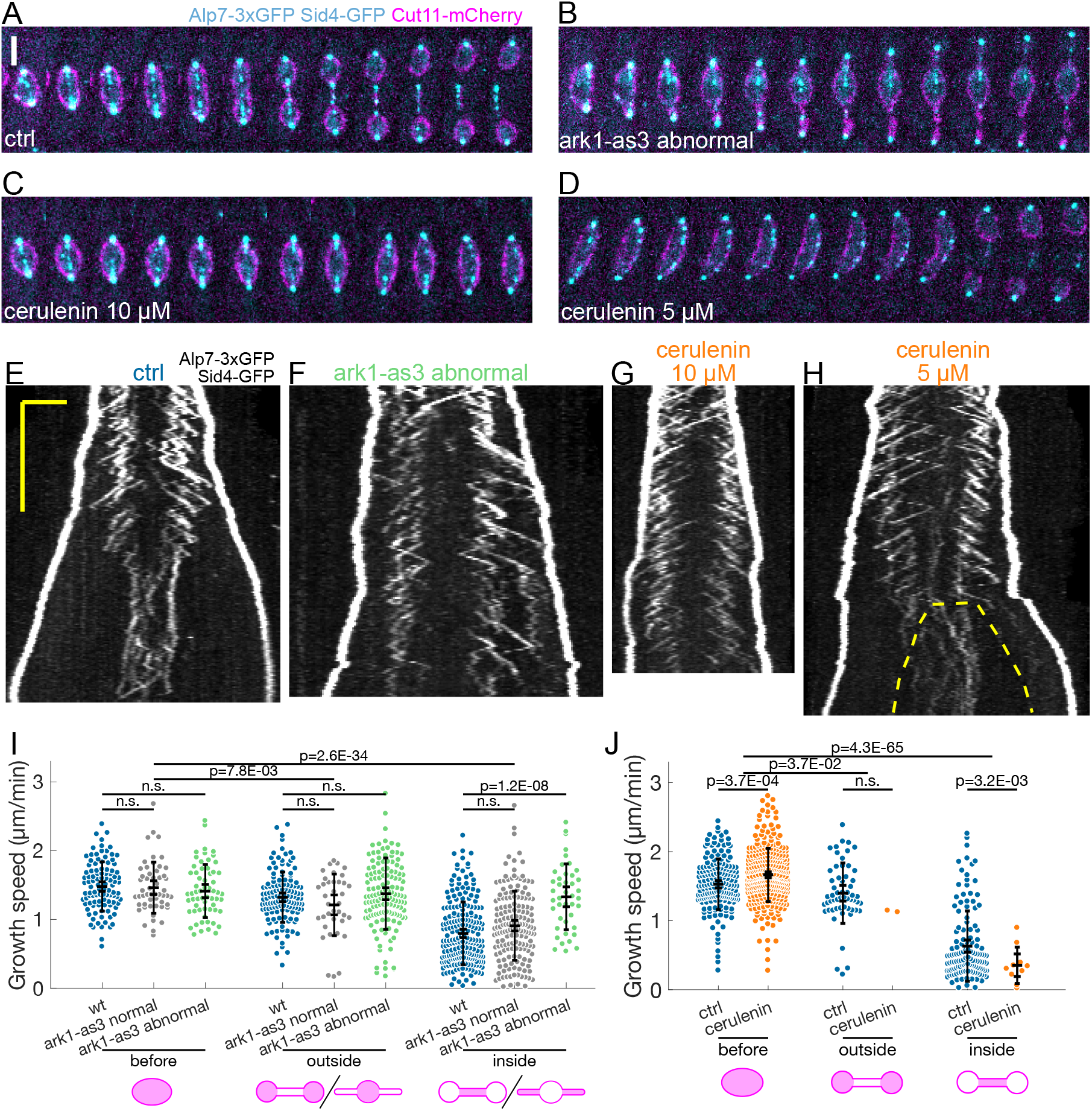
Preventing the dumbbell transition abolishes the switch from fast to slow microtubule growth. (**A-D**) Time-lapse images of mitotic spindles in cells expressing Alp7-3xGFP, Sid4-GFP (cyan) and Cut11-mCherry (magenta), in different conditions: (A) wild-type + DMSO (cerulenin control), (B) ark1-as3 with 5 μM 1NM-PP1, (C) wild-type with 10 μM cerulenin, (D) wild-type with 5 μM cerulenin in which the nuclear membrane eventually undergoes dumbbell transition. Time between images is 1 minute, scale bar 3 μm. (**E-H**) Kymographs of anaphase mitotic spindles in cells expressing Alp7-3xGFP, Sid4-GFP and Cut11-mCherry in the same conditions as (A-D). Time is in the vertical axis (scalebar 5 minutes), and space is in the horizontal axis (scalebar 2 μm). Dashed line in (H) outlines the nuclear membrane bridge formed after the dumbbell transition. See corresponding Cut11-mCherry kymographs in Fig. 4-Supplement 1A-D. (**I**) Microtubule growth speed in cells treated with 5 μM 1NM-PP1, blue: wild-type cells, grey: ark1-as3 cells that underwent normal chromosome segregation (‘ark1-as3 normal’), green: ark1-as3 cells that failed chromosome segregation and did not undergo dumbbell transition (‘ark1-as3 abnormal’). Events are categorised according to where the rescue occurred (see cartoons under x-axis, when two cartoons are drawn under a category, left is for wild-type and ‘ark1-as3 normal’, and right is for ‘ark1-as3 abnormal’). (**J**) Microtubule growth speed in wild-type cells treated with DMSO (blue) or 10 μM cerulenin (orange). Events are categorised according to where the rescue occurred (see cartoons below). Error bars represent 95% confidence interval of the mean and standard deviation. p-values are calculated with regression analysis, ‘n.s’ (not significant) indicates p>0.05. See Table S1. Number of events: (I) 446 (25 cells) wt, 257 (20 kymographs) ark1-as3 normal, 240 (17 cells) ark1-as3 abnormal, from 4 experiments. (J) 368 (28 cells) wt, 328 (23 cells) cerulenin, from 4 experiments.

Next, we examined cells treated with the fatty acid synthetase inhibitor cerulenin, which reduces cellular membrane availability, dramatically decreasing spindle elongation, and prevents dumbbell transition (Fig. 4C) [57]. Treating cells with 10 μM of cerulenin prevented dumbbell transition, and abolished the decrease in microtubule growth speed (Fig. 4C, G, Fig. 4-Supplement 1C). Furthermore, out of the 23 cells treated with cerulenin, we observed a clear bimodal growth velocity distribution only in 2 cells that underwent dumbbell transition despite the treatment (growth events in the ‘inside’ and ‘outside’ categories in Fig. 4J, and Fig. 4-Supplement 1E). By reducing cerulenin concentration to 5 μM, we observed two additional cells in which a reduction of microtubule growth speed after abrupt dumbbell transition was evident (Fig. 4H, Fig. 4-Supplement 1D, F).

In summary, our data shows that microtubule growth speed can remain unchanged for a period of time as long as wild-type anaphase B if the dumbbell transition is prevented (Fig. 4F, G). This suggests that the decrease in microtubule growth speed observed in wild type cells is not regulated by a “timer” mechanism, but instead occurs when the nuclear membrane bridge encloses the midzone.

### Ase1 is required for normal rescue distribution and for microtubule growth speed to decrease in anaphase B

Preventing the decrease in microtubule growth speed by cerulenin treatment or Aurora B inhibition did not compromise spindle stability nor overall microtubule organisation. This suggested that the way microtubule sliding and growth are coordinated is robust against deviations from the normal microtubule growth speed evolution. In fact, since rescues occur most often at midzone edges (Fig. 1G), microtubules are stable from pole to midzone, and only their distal parts including the plus ends alternate between assembly and disassembly phases. Therefore, promoting rescues at the midzone edges could be sufficient for the spindle to maintain a constant midzone length while sustaining sliding. We initially set out to test this mechanism experimentally by deleting ase1, a microtubule crosslinker that organises the midzone [8, 58] and recruits the rescue factor Cls1 [24]. As expected [8, 24, 58], in cells where ase1 was deleted the characteristic distribution of rescues was lost (Fig. 5B, D) and spindles collapsed due to loss of microtubule overlaps before reaching the typical final length (Fig. 5C, E, Fig. 5-Supplement 1A). Strikingly, we also observed that in ase1Δ cells microtubule growth velocity no longer decreased during anaphase B (Fig. 5F, Fig. 5-Supplement 1B). To better characterise the function of Ase1, we generated strains expressing mCherry-ase1 from the ase1 endogenous promoter, and cells overexpressing mCherry-ase1 from a P1nmt1 promoter (mCherry-ase1^OE^, Fig. 5-Supplement 1C-G), which resulted in a more than 10-fold increase of mCherry-Ase1 levels on the spindle (Fig. 5-Supplement 1H). Microtubule growth speed in mCherry-ase1^OE^ cells was similar to wild-type at early anaphase, but decreased more sharply with distance to the pole (Fig. 5-Supplement 1I). We next tried to reduce mCherry-ase1 levels by expressing mCherry-ase1 from a P81nmt1 promoter. However, the mCherry-Ase1 signal in these strains was undistinguishable from noise (data not shown). As an alternative, we quantified the levels of GFP-Ase1 in strains expressing GFP-ase1 either from its endogenous promoter, or from a P81nmt1 promoter (GFP-ase1^off^). We observed a decrease of approximately 3-fold in the GFP-Ase1 levels on the spindle in GFP-ase1 ^off^ cells compared to wild-type (Fig. 5-Supplement 1L, M). We then measured microtubule growth velocity in cells where expression of unlabelled ase1 was driven by a P81nmt1 promoter (ase1^off^, Fig. 5-Supplement 1J, K). ase1^off^ cells exhibited a slight increase in microtubule growth with respect to wild-type (Fig. 5-Supplement 1N), but normalised growth speed as a function of distance from the plus end to the closest pole at rescue was similar to wild type (Fig. 5-Supplement 1O). These results are reminiscent of those obtained when altering Cls1 levels (Fig. 2, Fig. 2-Supplement 1): the change in microtubule growth speed occurs across a wide range of Ase1 levels, suggesting that even though Ase1 is required for microtubule growth speed to decrease during anaphase B, this is unlikely to be a direct effect. Instead, it could be that Ase1 is required for the nuclear membrane bridge to have an effect on microtubule growth (see Discussion).

**Fig. 5:**
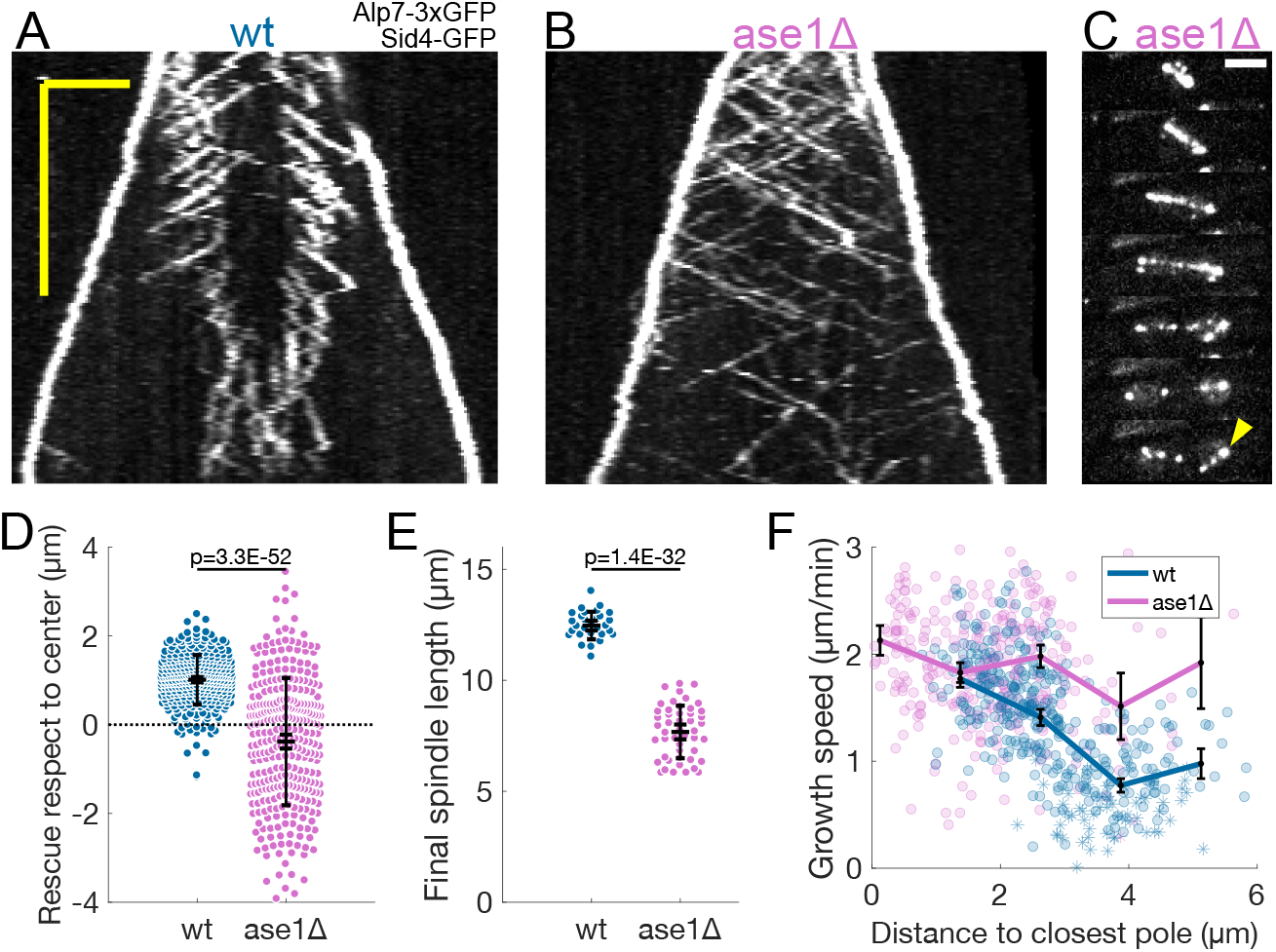
Ase1 is required for normal rescue distribution and for microtubule growth speed to decrease during anaphase B. (**A, B**) Kymographs of anaphase mitotic spindles in wild-type (A) and ase1Δ (B) cells expressing Alp7-3xGFP and Sid4-GFP. Time is in the vertical axis (scalebar 5 minutes), and space is in the horizontal axis (scalebar 2 μm). (**C**) Time-lapse images of a mitotic spindle in a cell where ase1 is deleted, expressing Alp7-3xGFP and Sid4-GFP. Time between images is 3 minutes, scale bar 3 μm. Arrowhead indicates spindle collapse that occurs due to loss of microtubule overlap. (**D**) Distribution of the position of rescues with respect to the spindle center in wild-type and ase1Δ cells. Dotted line marks the spindle center. (**E**) Distribution of final spindle length in wild-type and ase1Δ cells. Same data as in Fig. 5-Supplement 1A. (**F**) Microtubule growth speed as a function of the distance between the plus end and the closest pole at rescue (or first point if rescue could not be exactly determined) in wild-type (blue) and ase1Δ (pink) cells. Microtubule growth events of clear duration are shown as round dots, others as stars. Thick lines represent average of binned data, error bars in (F) represent 95% confidence interval of the mean. In (D, E) error bars represent 95% confidence interval of the mean and standard deviation. p-values correspond to T-test. Number of events shown: (E) 30 wt, 48 ase1Δ cells from 3 independent experiments. (D, F) 402 (34 cells) wt, 316 (39 cells) ase1Δ microtubule growth events from 4 independent experiments

In summary, Ase1 is required for rescue organisation and for microtubule growth speed to decrease during anaphase B.

### Promoting microtubule rescues at the midzone edge is sufficient to coordinate sliding and growth across a wide range of microtubule growth speeds

We next tested whether restricting rescues to the midzone edges is sufficient to coordinate microtubule sliding and growth. We developed a minimal mathematical model (see Methods) to compare scenarios that differ in how rescue is promoted inside the midzone. In our model, the spindle is initially composed of 9 antiparallel microtubules (Fig. 6A) that undergo dynamic instability. Based on the fact that Cls1 is recruited by Ase1 [24] and that Cls1 levels remain constant during anaphase (Fig. 2-Supplement 1B), we assume that a fixed amount of rescue factor (*R*) is distributed along the overlap between microtubules in a midzone of constant length. The rescue activity is either uniform along the midzone, or distributed according to a beta distribution with parameters *α* and *β*. Increasing *α* localises the rescue activity at the midzone edges to higher degrees, while keeping the total rescue factor constant (Fig. 6A). All the model parameters were derived from experimental measurements, except for *R, α* and *β* (Table S2). *R* was scanned and tuned to fit the experiments (see below). We explored *α*=*β*=1, corresponding to a uniform distribution, and *α* = 4, 8 and 12 with *β*=2, representing increasingly skewed distributions towards the midzone edge (Fig. 6A). We initially set the microtubule growth velocity to 1.6 μm/min (early anaphase speed, Fig. 1F), and aimed to reproduce the experimental distribution of positions of rescue and catastrophe at early anaphase (spindle length < 6 μm, Fig. 6B), measured from kymographs (Fig. 1C), and the total tubulin intensity as a function of spindle length (Fig. 6C), measured in cells expressing fluorescent *α*-tubulin (Fig. 1-Supplement 1E). Fluorescent tubulin intensity is proportional to polymerised tubulin [40, 59], so it can be compared to the total polymerised tubulin in a simulation by using a scaling factor (Fig. 6C, see Methods).

**Fig. 6:**
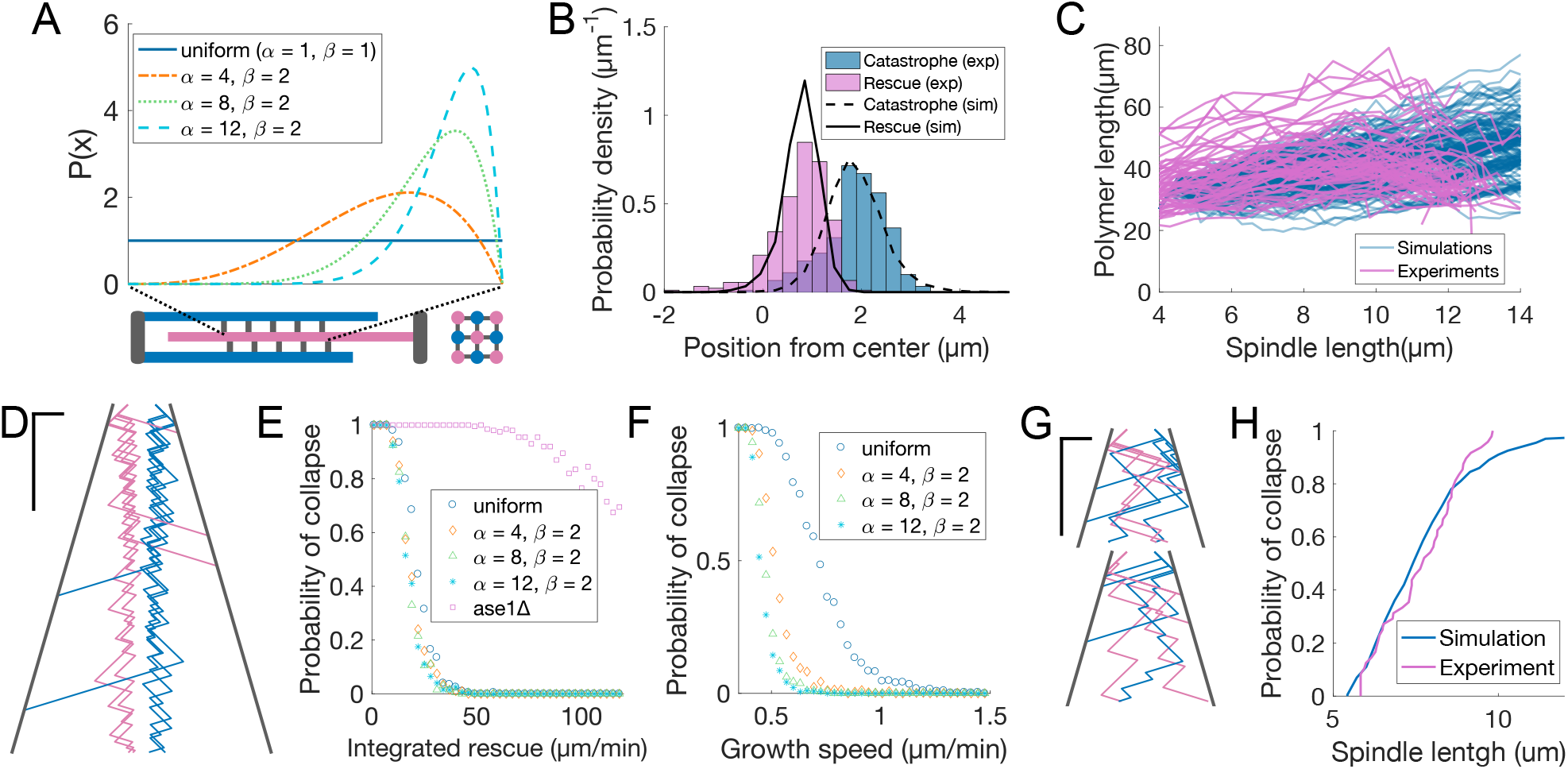
Promoting microtubule rescues at the midzone edge is sufficient to coordinate sliding and growth. (**A**) Arrangement of microtubules in simulations, shown as longitudinal (bottom left) and perpendicular (bottom right) sections of the spindle. Microtubules are colour coded according to their orientation, SPBs and connections between microtubules are shown in grey. The dashed lines linking the midzone edges to the extremes of the x-axis represent the fact that the parameter *x* maps the position along the midzone to a value that goes from zero to one. Curves inside the graph represent the value of *P*(*x*) from Equation 5 for parameters indicated in the legend. (**B**) Distribution of positions of microtubule catastrophe and rescue with respect to the spindle center in experiments (histograms, same data as Fig. 1G), and 200 simulations (lines), for *R* = 55μm/min, *α*=4, *β*=2. (**C**) Total polymerised tubulin as a function of spindle length in 200 simulations (blue) and total mCherry-atb2 intensity (scaled) as a function of spindle length in 60 cells from 6 independent experiments (pink). Simulation Parameters as in (B). (**D**) Kymograph generated from a simulation with parameters as in (B). Plus-ends are colour coded according to their orientation, SPBs are shown in grey. Time is in the vertical axis (scalebar 5 minutes), and space is in the horizontal axis (scalebar 2 μm). Total simulated time is 20 minutes. (**E**) Probability of spindle collapse as a function of the total rescue factor (*R*). Each dot represents a set of 200 simulations with equal parameters (Table S2), and its value on the y axis is the fraction of the simulations in which the spindle collapsed in the 20 minutes of simulated time. (**F**) Same as (E), but each dot represents 500 simulations, for various values microtubule growth speed. (**G**) Simulation kymograph as in (D), for ase1Δ, where rescue activity is uniformly distributed along the whole length of microtubules. Both simulations ended with a spindle collapse due to loss of the overlap between antiparallel microtubules. *R* = 34 μm/min (**H**) Cumulative distribution of final spindle length in ase1Δ cells (pink, same data as Fig. 5E), and in 200 ase1Δ simulations (blue). See Table S2 for simulation parameters.

A good agreement with the experimental data was obtained with *α*=4, *R*=55 μm/min (Fig. 6B-D, Fig. 6-Supplement 1D). For higher values of *R*, the spindle was stable (Fig. 6E) but, due to the higher microtubule stability, the total polymerised tubulin increased steadily, unlike in the experiment, where it remained more or less constant throughout anaphase (Fig. 6C). Importantly, the distributions of rescues and catastrophes were similar to the experimentally observed ones when the rescue factor was uniformly distributed along the midzone (Fig. 6-Supplement 1E) because even in that case, the most likely place to have a rescue is the first position where it can happen.

For microtubule growth speed characteristic of early anaphase (1.6 μm/min), the spindle stability was not affected by the distribution of rescue rate within the midzone (Fig. 6E). In contrast, if we kept *R* as in Fig. 6B-D, but decreased the microtubule growth speed, we observed that spindles with skewed rescue distributions were more stable (Fig. 6F). When growth velocity is low, a rescued microtubule might not exit the midzone before undergoing catastrophe, and therefore will “miss” a fraction of the midzone where it could have been rescued. Increasing the rescue rate close to the midzone edge makes it more likely that plus ends of rescued microtubules are outside the midzone by the time they undergo catastrophe, which increases their chances to be rescued again.

In summary, this simple model shows that promoting rescues at midzone edges is sufficient to maintain the microtubule overlap and sustain sliding across a wide range of microtubule growth speeds. Within our limited exploration, it suggests that accumulating the rescue activity at the midzone edges might represent an optimal localisation of this activity, in the context of anaphase B.

### Loss of microtubule rescue organisation leads to spindle collapse

Finally, we tested whether the loss of rescue organisation observed in ase1Δ cells (Fig. 5D) could be partly responsible for spindle collapse (Fig. 5C, E). We modified the model so that the total amount of rescue factor was distributed all along microtubules, and the rescue rate was the same anywhere on the spindle. By scanning the parameter that represents the total rescue factor (*R*), we found that most spindles collapsed due to loss of antiparallel overlap of microtubules in 20 minutes of simulated time, even for values of *R* way higher than those required to maintain stability in spindles with midzone (Fig. 6E). This is similar to what happens in ase1Δ cells, where interpolar microtubules coming from opposite poles often lose their connection prior to reaching the typical final spindle length (Fig. 5C, E). For *R*=34 μm/min, this modified model closely reproduced the distribution of spindle length at collapse observed in ase1Δ cells (Fig. 6G, H, Fig. 6-Supplement 1F), suggesting that in these cells a combination of lower rescue rate and loss of rescue organisation leads to spindles collapsing prior to reaching the typical final spindle length.

## Discussion

We have measured microtubule dynamics during anaphase B in *S. pombe* to examine how microtubule polymerisation and sliding are coordinated. We found that: (1) Wrapping of the nuclear membrane around the spindle midzone reduces microtubule growth speed. (2) Rescues occur preferentially at midzone edges. We then developed a model showing that organising rescues in this manner is sufficient to coordinate microtubule growth and sliding across the wide range of microtubule growth speeds observed experimentally.

### Wrapping of the nuclear membrane around the spindle midzone reduces microtubule growth speed

In higher eukaryotes, microtubule dynamics are increasingly suppressed during anaphase B by kinesin-4 [11, 36, 60]. We have found that in *S. pombe*, which lacks kinesin-4, microtubule growth speed also decreases during anaphase B (Fig. 1E). Microtubules exist in two states (fast and slow growing) characteristic of early and late anaphase (Fig. 1F). The transition between these states occurs independently of several motors and MAPs that regulate microtubule dynamics (Fig. 2), and happens when plus ends enter the nuclear membrane bridge formed after the dumbbell transition (Fig. 3). Furthermore, microtubules do not switch to the slow growing state if bridge formation is prevented by cerulenin treatment or Aurora B inhibition (Fig. 4). Our data suggests that microtubule growth speed is mainly governed by spatial cues, rather than a “timer” (Fig. 2K, Fig. 2-Supplement 1J), which is reminiscent of recent studies showing that late mitotic events respond to spindle length and not time [61, 62].

It is tempting to speculate on mechanisms that could decrease microtubule growth speed when plus ends enter the nuclear membrane bridge. A protein present at the bridge might directly reduce microtubule growth speed. Alternatively, close contact of microtubules with the nuclear membrane or its associated proteins might physically hinder growth [43, 63]. Finally, since there is no diffusion between the nuclear membrane bridge and the daughter nuclei [28, 30], the availability of tubulin dimers or another factor could limit microtubule growth when plus ends enter the bridge. All these possibilities are compatible with the fact that Ase1 is required for microtubule growth speed to decrease during anaphase (Fig. 5F), as previous reports suggest that Ase1 is involved in the interaction of the nuclear bridge with the spindle: in *S. cerevisiae* the nuclear bridge acts as a diffusion barrier between daughter nuclei only in the presence of Ase1 [30], and in *S. pombe* Ase1 is required for the local concentration of nuclear pore complexes at the nuclear membrane bridge [29]. It would be interesting to test whether microtubule growth speed also decreases during anaphase in the closely related fission yeast *S. japonicus*, which does not form a nuclear membrane bridge during anaphase [57]. Importantly, membranes also tightly wrap around the spindle in other systems, such as the midbody in animal cells [64], or the phragmoplast in plants [65], and may affect microtubule dynamics.

### Promoting microtubule rescues at the midzone edges grants coordination of sliding and growth

The microtubule rescue factor Cls1 (CLASP) is recruited to the midzone by the crosslinker Ase1 (PRC1) in several species [24–26], and it had been previously proposed that this would restrict rescues to the midzone [24]. Indeed, we observed that rescues occur most frequently at midzone edges (Fig. 1G). Thus, spindle microtubules remain stable from the pole to the midzone edge, and only their distal parts including the plus ends alternate between assembly and disassembly (Fig. 6D). Simulations show that promoting microtubule rescue at midzone edges is sufficient to maintain the overlap between microtubules and sustain sliding across a wide range of microtubule growth speeds (Fig. 6F), indicating that this mechanism is robust against perturbations on microtubule growth. Our study suggests that *S. pombe* cells adopted this mechanism to ensure spindle integrity, rather than a precise regulation of microtubule growth. This is supported by the fact that deletion of Ase1 [8, 58] and inactivation of Cls1 [24] lead to spindle collapse, while perturbing the normal microtubule growth speed evolution during anaphase B has no effect on spindle stability or organisation (Fig. 4). In higher eukaryotes the situation is likely more complex: in HeLa cells, depletion of kinesin-4 leads to highly dynamic interpolar microtubules [11], but spindles still elongate with normal speed during anaphase B [4], indicating that a precise regulation of microtubule growth is not required. However, depletion of PRC1/Ase1 does not perturb anaphase spindle elongation either [4], suggesting that if microtubule rescue organisation is required for spindle stability, it may rely on a mechanism other than recruitment of CLASP by PRC1. It would be important to understand the relative contribution to spindle stability of these mechanisms, and others not present in yeast during anaphase B, like microtubule nucleation [21, 22].

We do not know the molecular mechanism by which rescues happen more often at midzone edges. We can speculate that this might be related to the fact that protofilaments curl outwards when microtubules shrink [66]. Indeed, the distances between adjacent microtubules in the midzone are comparable to the radius of curling protofilaments [40, 66], such that steric collisions alone could promote protofilament straightening, enhancing rescue similarly to what has been proposed for kinetochore protein Dam1 [66, 67]. Recent *in vitro* studies have combined PRC1, CLASP and dynamic microtubules [14, 68], so it would be interesting to see whether in a reconstitution resembling the yeast spindle rescues also happen at the edges, which would indicate that this is an inherent property of PRC1/CLASP systems.

### Consequences of keeping a fixed amount of rescue factor in the midzone

Following our observation that Cls1 levels on the spindle remain constant during anaphase (Fig. 2-Supplement 1B), we have assumed that a fixed amount of rescue factor distributes along the midzone overlap, which makes spindles increasingly stable as they lose microtubules. This mechanism might operate in cells, since anaphase spindles can last for up to 40 minutes if spindle disassembly is delayed (Fig. 2-Supplement 1I). Additionally, assuming that a fixed amount of rescue factor distributes along overlaps makes rescue rate proportional to the number of neighbours of a microtubule. Therefore, when a mi-crotubule fully depolymerises, its neighbours oriented in the opposite direction have then one fewer neighbour, while the microtubules in the same orientation as the lost one keep the same number of neighbours. This simple effect promotes even loss of microtubules from both poles, which is necessary to prevent spindle collapse during anaphase B in *S. pombe*. To illustrate the need for even loss of microtubules from both poles, let us consider a representative spindle that has 9 microtubules at anaphase B onset. By the end of anaphase, approximately 5 microtubules will remain [40]. Thus, in the absence of a mechanism driving even loss of microtubules, all microtubules would have a probability ~1/2 of being lost. The probability of losing all microtubules from either pole would then be 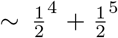, so approximately 10% of spindles would collapse because of this. In contrast, spindles do not collapse in our simulations, where this mechanism operates. In 500 simulations with parameters as in Fig. 6B, C, where spindles start anaphase with 9 microtubules, and have on average 5 microtubules after 15 minutes (the typical duration of anaphase B) we observed no spindle collapse (Fig. 6F).

### Cross-talk between microtubule sliding and growth

We saw no evidence of cross-talk between microtubule sliding and growth. Klp9 has no noticeable effect on microtubule growth speed, despite its growth promoting activity in monopolar spindles [17]. We have not seen growth-limited sliding in unperturbed spindles either: although microtubule growth speed decreases by ~60% (Fig. 1F), spindle elongation speed remains constant throughout anaphase B (Fig. 1-Supplement 1A). In *S. cerevisiae* and *Drosophila*, deletion or depletion of Kinesin-8, which decreases catastrophe rate [15, 69], increases sliding speed and final spindle length [15, 69]. This could indicate that sliding is growth-limited in these organisms. On the other hand, in *S. cerevisiae*, deletion of kinesin-8 kip3 also delays spindle disassembly [70]. Other mutations that delay spindle disassembly (like cdh1Δ, which prevents the degradation of midzone crosslinkers) produce hyperelongated spindles, similarly to kip3 deletion [70]. In both kip3Δ and cdh1Δ, spindle elongation only stops when spindles are cut by the cytokinetic ring [70], suggesting that no mechanism exists to stop sliding during anaphase B other than the disassembly of the midzone, and the same has been observed in *S. pombe* [28]. Growth-limited sliding does not seem to occur in HeLa cells either, as depletion of kinesin-4 does not affect spindle elongation velocity [4]. Ultimately, measuring microtubule dynamics in these systems will reveal their impact on microtubule sliding. Nevertheless, growth-limited sliding is widely conserved, and may contribute to spindle stability in *S. pombe* when microtubule dynamics are perturbed, such as during starving [71].

## Materials and methods

### Production of S. pombe mutant strains

All used strains are isogenic to wild-type 972 and were obtained from genetic crosses, selected by random spore germination and replicated on plates with corresponding drugs or supplements. Gene deletion and tagging was performed as described previously [72], following [73]. All strains, oligonucleotides and plasmids used are listed in the supplementary file ‘strains_plasmids_oligos.xlsx’.

### Fission yeast culture

All *S. pombe* strains were maintained at 25° in YE5S plates and refreshed every third day. One day before the microscopy experiments, cells were transferred to liquid YE5S culture, and imaged the next day at exponential growth. For all experiments except for Fig. 3, the cells were grown overnight in YE5S liquid medium at 25°. For experiments in the absence of thiamine (used to induce overexpression of klp9, Fig. 3), cells were pre-grown in liquid YE5S, then washed three times with deionised water, transferred to EMM supplemented with adenine, leucine and uracil, and incubated 18-22 hours at 25° prior to the microscopy experiment. For cerulenin treatment, cells were incubated for 1 hour in liquid medium with the indicated cerulenin concentration prior to the experiment (Sigma-Aldrich, stock solution was 10 mM in DMSO), the same volume of DMSO was added in control. For Aurora B inactivation, 5 μM of 1NM-PP1 was added to both wildtype and ark1-as3 cells (Sigma-Aldrich, stock solution was 5uM in DMSO), and cells were imaged immediately.

### Live-cell microscopy

For live-cell imaging, cells were mounted on YE5S agarose pads, containing 4% agarose [74]. For the experiments in Fig. 4, the drug or DMSO was added at the same concentration in the agarose pad as in the liquid medium.

Imaging of data from figures 1, 1-Supplement 1, 1-Supplement 2, 2 (mal3, cls1, klp9 and their corresponding wt) and 5-Supplement 1 (ase1^off^ and its corresponding wt) was performed at 27° with an inverted Spinning disk confocal microscope Eclipse Ti-E (Nikon) with Spinning disk CSU-X1 (Yokogawa), equipped with Plan Apochromat 100/1.4 NA objective lens (Nikon), a PIFOC module (perfect image focus), and sCMOS camera Prime 95B (Photometrics), integrated in Metamorph software by Gataca Systems. For GFP and mCherry imaging, lasers of 488 nm (100 mW) and 561 nm (50 mW) were used. Movies used for kymographs were imaged using a Live-SR module (Gataca systems).

The rest of the imaging was performed at 27° with an inverted Spinning disk confocal microscope Eclipse Ti-2 (Nikon) with Spinning disk CS-W1 (Yokogawa), equipped with objective 100x CFI Apo VC/1.4 NA (Nikon), a PIFOC module (perfect image focus), and sCMOS camera Prime 95B (Photometrics), integrated in Metamorph software by Gataca Systems. For GFP and mCherry imaging, lasers of 488 nm (150 mW) and 561 nm (150 mW) were used. Movies used for kymographs were imaged using a Live-SR module (Gataca systems).

Movies for kymographs were acquired as follows: in the GFP channel, images were acquired as stacks of 5 planes spaced 0.5 μm without binning every 4 seconds during 15 minutes (except for mCherry-Ase1 movies, where the images were acquired every 5 seconds). Exposure was 100ms (Gain 3). In the mCherry channel, for Cut11-mCherry images were acquired as single stacks, every 16 seconds without binning during 15 minutes. For mCherry-ase1, images were acquired as stacks of 5 planes spaced 0.5 μm without binning every 8 seconds during 15 minutes. Exposure was 200ms (Gain 3). Movies where intensity measurements were made were acquired as stacks of 11 planes spaced 0.5 μm without binning every minute during 90 minutes. Exposure was 100 ms (Gain 3) in all channels. The remaining movies were acquired as stacks of 7 planes spaced 1 μm without binning every minute during 90 minutes. Exposure was 100 ms (Gain 3) in all channels.

To minimise the inter-experiment variance, all the data shown within the same plot was acquired in parallel. In the case of movies used for kymographs this meant that samples of the conditions tested were alternated on the microscope. In the case of one minute interval movies, positions corresponding to all conditions were imaged simultaneously.

### Statistical analysis

All statistical analysis was performed in Matlab. Student t-test was used to compare univariate distributions in Fig. 5, 2-Supplement 1A and 3-Supplement 1. Regression analysis and 2-way anova test were used to compare the conditions in Fig. 3 and 4 (see Table S1).

### Image and data analysis

All feature detections, kymographs and length measurements were done in maximal projections of fluorescence microscopy images. Intensity measurements were performed on summed projections.

#### Spindle detection

Sid4-GFP Alp7-3xGFP and fluorescence tubulin movies were pre-processed with the Fiji distribution of Imagej (https://imagej.net/Fiji). The plugin *Trainable Weka Segmentation* [75] was used to generate probability images of each frame in maximal projections. Each probability image has the same size as the original maximal projection, with values in [0,1]. Higher values correspond to higher likelihood of that pixel corresponding to a spindle. The probability images were later used with a custom Matlab script to detect spindles, which is publicly available (link). The user is required to draw the profile of each analysed cell by hand. Since generally cells do not move between frames, the same profile can be used for all the frames, but it can be changed between frames if necessary.

To find the spindles, we use the pixels inside the cell mask in probability images for which the probability is bigger than 0.8 (This gives a good segmentation of the spindle). Pixels are defined by position (*X_i_*, *Y_i_*) and probability (*P_i_*). We find the parameters (*X*_0_,*Y*_0_,*θ*,*α*) that maximise a functional *F* defined as:

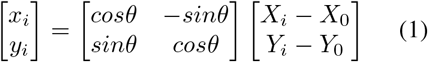

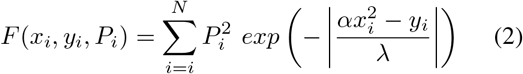

Equation 1 rotates a point by angle *θ* around (*X*_0_, *Y*_0_). Equation 2 defines an objective function, aiming to fit a spindle defined by *g*(*x*) = *αx*^2^, a polynomial of degree 2 in the rotated axis. F is a weighted sum computed from the probability *P_i_* and exponential terms derived from the distance to *g*(*x*), with a characteristic width λ=0.55 μm. To find the polynomial that best matches the spindle, we find the values of (*X*_0_,*Y*_0_,*θ*,*α*) that maximise *F*. Then, to find the spindle edges, we project all the points of the probability image on the polynomial *g*(*x*). The edges are defined as the two points on *g*(*x*) that contain all projections with *P_i_* >0.6 between them. We define the length of the spindle as the length of the curve *g*(*x*) between these two edges. We fit data in each frame to this function, starting by the first time point (Movie 1). Initially, we set *α* to zero, until we find a spindle of length greater than 6 μm, since shorter spindles are straight and well described by a line. If the algorithm fails to find the trace of the spindle, the user can draw the spindle by hand. In Alp7-3xGFP Sid4-GFP movies used for kymographs we use a slight variation in which first the SPBs are detected, since Sid4-GFP signal is stronger, and we constrain the fit to polynomials passing by those two points.

#### Intensity measurements

To measure the intensity of fluorescent tubulin and MAPs, we used intensity profiles (Matlab *improfile* function) on summed projections along curves obtained with the spindle detection algorithm described above. The curve points were evenly spaced at a distance of 1 pixel. We used 7 parallel curves, separated by 1 pixel, the central one coinciding with the orthogonal polynomial *g*(*x*). The total width corresponds to 0.77 μm in our microscope. We consider this to be the signal region. As background, we subtracted the median of two parallel regions of 4 pixels (0.44 μm) width flanking the signal region. The intensity was the sum of the intensities of all the points in the signal region after background subtraction. For the density of Cls1-3xGFP (Fig. 2-Supplement 1C, E), we only measured intensities of Cls1-3xGFP and mCherry-Atb2 in the central 2 μm of the spindle.

#### Kymograph analysis

To generate kymographs, we used intensity profiles (Matlab *improfile* function) on maximal projections of Sid4-GFP Alp7-3xGFP movies on which a Gaussian filter of 1 pixel width was applied, along curves obtained with the spindle detection algorithm described above. For each of the frames in the movie, we constructed a maximal projection along a curve calculated from the fit, where points were evenly spaced at a distance of 1 pixel. We projected all the points that were at a distance of 3 or less pixels from the spindle trace, and this yielded a vector of intensities for each time point. To align the vectors into a kymograph we used the center of mass of the maximal projection of the movie in time as a reference, and we aligned all the vectors such that this point would be at the center of the kymograph. Cut11-mCherry and mCherry-Ase1 kymographs were constructed the same way, but no Gaussian filter was applied.

To annotate the kymographs, we developed a Matlab program, KymoAnalyzer, (link) allowing one to draw polygonal lines to mark the poles of the spindle and the microtubule growth events in the Sid4-GFP Alp7-3xGFP channel, and the membrane bridge edges in the Cut11-mCherry channel. The manually drawn lines are then resampled by interpolating the position at every time point. Duration of the growth event was defined as the time between apparition of the comet and its disappearance. To calculate the microtubule growth speed, the length vs. time curve was fitted to a first order polynomial using Matlab *polyfit*. The spindle center is defined as the middle between the poles. Kymographs of Cut11-mCherry were also used to determine the spindle length at which dumbbell transition occurred (Fig. 3-Supplement 1D).

#### Anaphase onset determination

To determine the onset of anaphase, we fitted the curves of length of spindle in time to the function *G*(*t*), with fitting parameters (*L*_0_, *t*_1_, *t*_2_, *s*_1_, *s*_2_, *s*_3_):

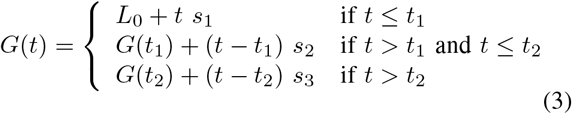

*G*(*t*) is a series of three linear fits that start at *L*_0_ at *t* = 0. Anaphase onset is determined by *t*_2_, and we use *s*_3_ as the anaphase B spindle elongation velocity. In general, fits correctly identified the switch in velocity from Phase II to III seen in Fig. 1-Supplement 1A. In some cases, particularly in klp9Δ, where the anaphase velocity decreases in time, *t*_2_ was fixed manually if the fit gave an obviously wrong result.

#### Inference of anaphase time from spindle length

In Fig. 2-Supplement 1J and Fig. 3-Supplement 1E we infer anaphase timing from spindle length. To do this, we use the curves of spindle length in time obtained in movies in which images were acquired every minute. We fit the curves of spindle length in time during anaphase to first order polynomials using Matlab *polyfit* (Fig. 2-Supplement 1K, Fig. 3-Supplement 1H), with anaphase onset determined as indicated above. Using the fit parameters, we then transform the spindle length measured in kymographs into time.

#### Public Matlab libraries used in this study

- geom2d - 2D geometry library
- hline-vline - plotting tool
- UnivarScatter - univariate scatter plots

### Computational model

The simulation is written in Python and available under an open source license https://github.com/manulera/simulationsLeraRamirez2021.

All microtubules are straight and aligned with the X axis, with the origin corresponding to the spindle center. The spindle poles, which contain the microtubule minus ends, are positioned symmetrically around the origin. Microtubules growing from one pole are orientated towards the opposite pole (Fig. 6A). Their positions in the YZ plane are recorded by an index representing their position on a transverse chequerboard lattice (Fig. 6-Supplement 1A). This index may change when the spindle “reorganises” (Fig. 6-Supplement 1A, and see below). For simplicity, we assume that the midzone is centered and has a constant length (*L_m_*), bypassing the complex dependence of midzone on MAPs binding to microtubule overlaps. In each simulation, *L_m_* is drawn randomly, sampling from a fit of the midzone length data shown in Fig. 1G to a normal distribution (Fig. 6-Supplement 1C). Each microtubule has its own length, and can exist in a growing or shrinking state. Microtubule growth, shrinkage and sliding occur at constant speeds (*υ_p_, υ_d_, υ_s_* Table S2). The duration of microtubule growth events is sampled from a fit of the cumulative distribution of duration of microtubule growth events shown in Fig. 1-Supplement 1C to:

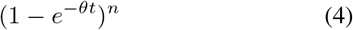

This function represents the probability of *n* events occurring in an interval of time *t*, if they are independent and occur with a constant rate *θ*. For *θ*=3.17 min^-1^ and *n*=8.53 (see Table S2), we obtained a good agreement with the data (Fig. 6-Supplement 1D). This suggests that catastrophe is not a single step process, agreeing with what was proposed in [48]. We assume for simplicity that a growing microtubule reaching the spindle pole immediately undergoes catastrophe. Shrinking microtubules may be rescued if their plus end is inside the midzone. Based on the fact that Cls1 is recruited by Ase1 [24] and that Cls1 levels remain constant during anaphase (Fig. 2-Supplement 1B), we assume that a fixed amount of rescue factor is distributed along the overlap between microtubules. Therefore, the integral of rescue rate (*r*) along the overlap length of the midzone is constant, and given by 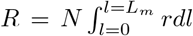, where N is the number of microtubule pairs in the square lattice. The rescue rate of a microtubule with *n* neighbours is given by the following expression:

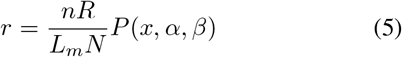

*P*(*x, α, β*) is the probability density of a beta distribution with parameters *α* and *β* as a function of *x*, a reduced coordinate that maps the position of microtubule plus ends in the midzone from zero to one, with zero corresponding to the midzone edge closest to the spindle pole (Fig. 6A). The beta distribution offers differently shaped distributions of rescue activity while keeping *R* constant. The spindle can reorganise when a microtubule is lost by complete depolymerisation. We assume that another microtubule can rearrange in the YZ plane to maximise antiparallel connections, as these are the arrangements observed by electron microscopy [40, 41]. Microtubules move to another position of the square lattice, of the same orientation, to increase the compactness of the structure. This can happen in two cases, as shown in Fig. 1-Supplement 1A. (1) A microtubule is lost and another one moves in its place (2) A neighbour of the lost microtubule moves to another available position. The moves only occur if they increase the number of antiparallel connections.

Simulations are initialised with a 4 μm long spindle, with 9 microtubules of 3 μm of length arranged in a square lattice as shown in Fig. 6A. Initially they all start in a growing state. The system is evolved using a time step *h* of 0.01 min. The new positions of spindle poles are calculated as *x_t+h_* = *x_t_* + *υ_s_h*, and microtubule lengths are updated similarly. Catastrophe and rescue are stochastic events simulated using the kinetic Monte Carlo method. At the beginning of the simulation, and whenever a microtubule rescue occurs, the time until the next catastrophe for a given microtubule (*τ_c_*) is sampled from the probability distribution in Equation 4. At every time point, *h* is subtracted from the *τ_c_* of each growing microtubule, until *τ_c_* is smaller than zero, at which point the microtubule switches to shrinkage. Shrinking microtubules can only be rescued inside the midzone, and at every time point the rescue rate (*r*) is given by Equation 5. The probability to be rescued during *h*, 1 – *e*^−rh^ is tested against a random number, uniformly distributed in [0,1]. At every time point we check if the spindle is “broken” by testing whether the longest microtubules of each pole are too short to overlap. The simulation finishes if the spindle breaks or when it reaches 20 minutes of simulated time.

The simulations of ase1 spindles are performed similarly, but the rescue rate is the same anywhere on the spindle and is given by *r* = *R*/2*L_t_*, where *L_t_* is the total length of microtubules. In this model, we assume that a fixed amount of rescue factor is distributed all along microtubules and not along the overlap between neighbours. The factor 2 in the denominator ensures consistency between both models. The fitting parameters for Equation 4 in these simulations are *θ*=6.8 min^-1^ and *n*=2.5 (Table S2).

#### Comparison of simulations and experiments

To compare microtubule polymer length as a function of spindle length in simulations with total tubulin intensity as a function of spindle length in experiments (Fig. 6-Supplement 1D), we binned the data based on spindle length, in bins of 1 μm width. For each bin, we calculated the average value of polymerised tubulin in simulations (*s*) and tubulin intensity in experiments (*e*). Then we calculated a scaling factor (*f*) that minimised the difference between them, and used the sum of square differences across all *N* bins 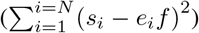 to score the similarity between simulations and experiments.

To compare the distribution of spindle length at collapse in ase1Δ simulations and experiments (Fig. 6-Supplement 1F), we calculated their empirical cumulative probability distributions with Matlab function *ecdf*. We then used the Kolmogorov-Smirnov distance between these distributions to score the similarity between simulations and experiments. The Kolmogorov-Smirnov distance is the supremum of the distances between cumulative probability distributions (*F*) across all their domain (*sup_x_* |*F*_1_(*x*) - *F*_2_(*x*)|).

## Acknowledgements

We thank Lara Katharina Krüger and Serge Dmitrieff for discussions and critical reading of the manuscript. We thank Stefania Castagnetti (LBDV, Sorbonne université), Jonathan Millar (University of Warwick), Ken Sawin (University of Edinburgh), Sylvie Tournier (Université de Toulouse) and Rafael R. Daga (universidad Pablo de Olavide, Sevilla) for kindly providing strains and plasmids used in this study. We thank Vincent Fraisier and Lucie Sengmanivong for the maintenance of microscopes at the PICT-IBiSA Imaging facility (Institut Curie), a member of the France-BioImaging national research infrastructure. We thank the Japan National BioResource Project Yeast Genetic Resource Center (Osaka City University, Osaka University and Hiroshima University) for providing strains.

## Additional information

### Author Contributions

M.L.R: Conceptualisation, Data curation, Formal analysis, Validation, Investigation, Visualisation, Methodology, Writing - original draft, Writing - review and editing, Software - development; F.J.N: Conceptualisation, Writing - review and editing, Supervision, Software - review; P.T.T: Conceptualisation, Writing - review and editing, Supervision, Funding acquisition, Project administration.

---

**Fig. 1-Supplement 1:**
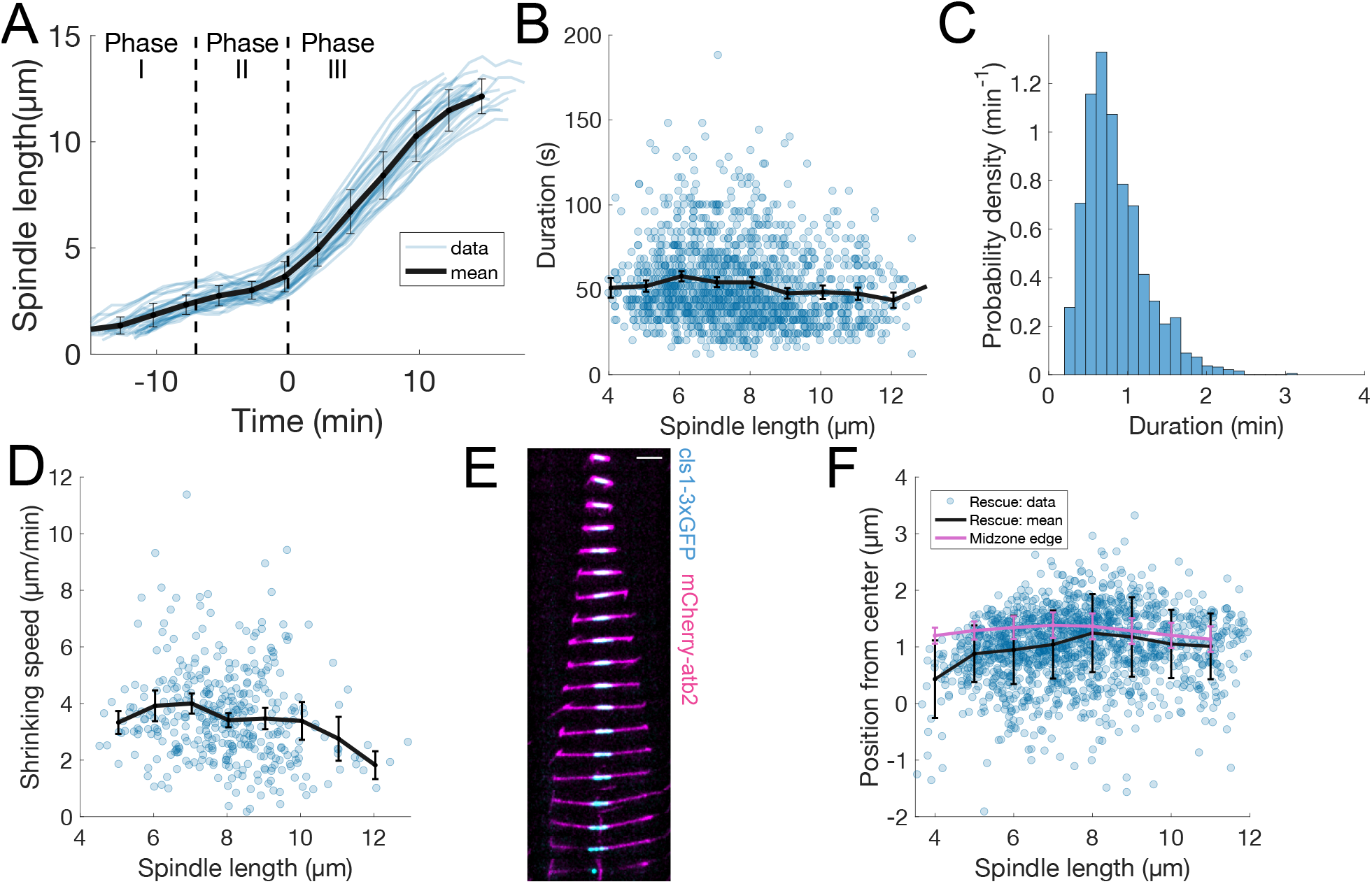
Characterization of microtubule dynamics during *S. pombe* anaphase. (**A**) Spindle length measured from Sid4-GFP Alp7-3xGFP signal in time (Fig. 1B), from spindle formation to spindle disassembly. Time is zero at anaphase onset. The dashed lines indicate transition between mitotic phases. Thin lines represent individual trajectories, thick black line represent average of binned data. Error bars represent 95% confidence interval of the mean. (**B**) Microtubule growth event duration as a function of spindle length at rescue. Thick black line represent average of binned data, error bars 95% confidence interval of the mean. (**C**) Histogram of values in (B). (**D**) Microtubule shrinkage speed as a function of spindle length at catastrophe. Thick black line represent average of binned data, error bars 95% confidence interval of the mean. (**E**) Time-lapse images of the anaphase spindle of a cell expressing mCherry-Atb2 (magenta) and Cls1-3xGFP (cyan). Time between images is 1 minute, scale bar 3 μm. (**F**) Position of rescues with respect to spindle center in cells expressing Sid4-GFP and Alp7-3xGFP (dots and black line), and position of midzone edge with respect to spindle center in cells expressing mCherry-Atb2 and Cls1-3xGFP (pink line). Thick black line represent average of binned data, error bars standard deviation of binned data. Number of observations: (A) 30 cells, from 2 independent experiments, (B, C, F) 1425 microtubule growth events, from 119 cells, from 13 independent experiments (D) 391 microtubule shrinkage events, from 89 cells, from 13 independent experiments (F) 60 cells, from 6 independent experiments.

**Fig. 1-Supplement 2:**
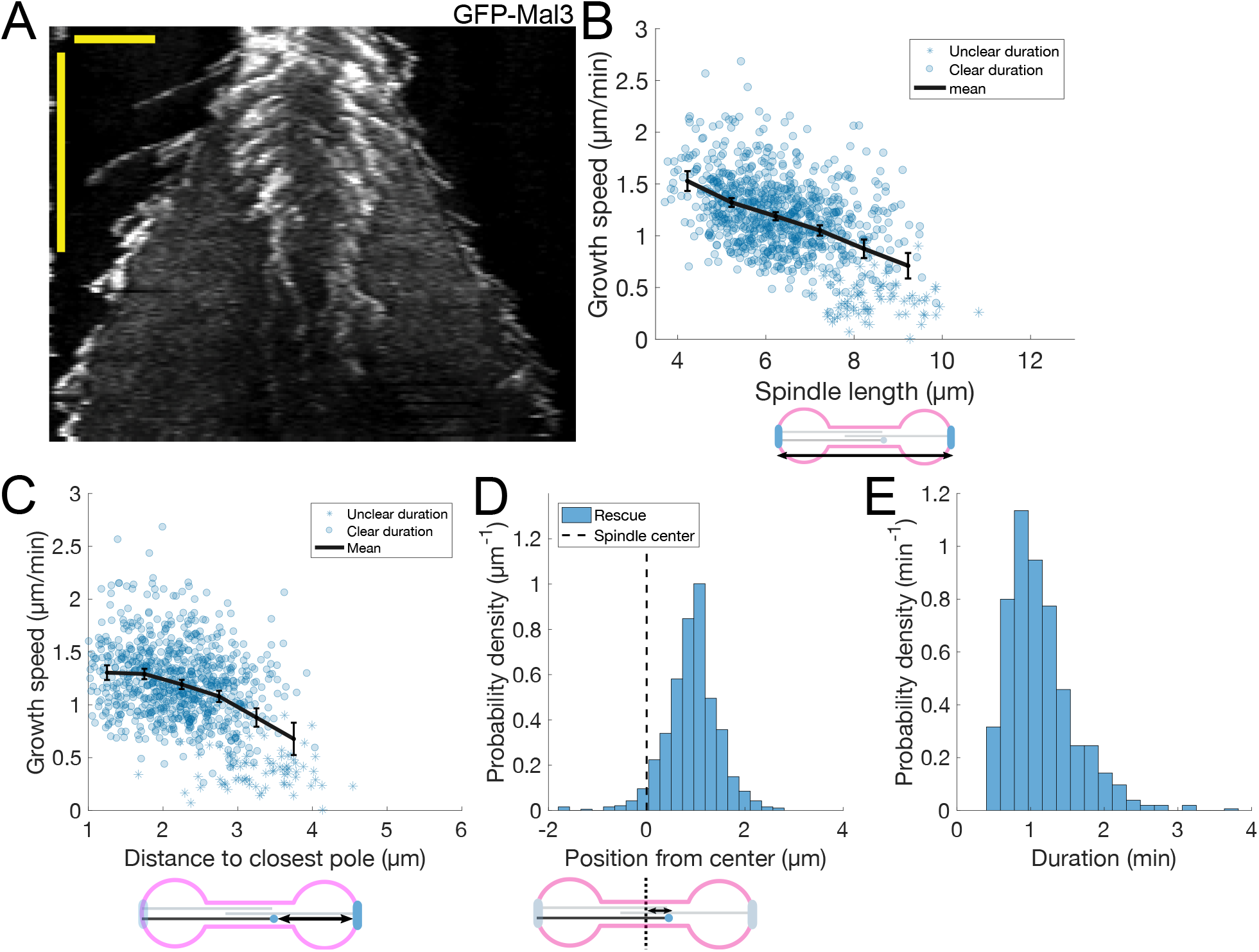
Microtubule dynamics during anaphase in *S. pombe* cells expressing GFP-Mal3. (**A**) Kymograph of a mitotic spindle during anaphase of a cell expressing GFP-Mal3. Time is in the vertical axis (scalebar 5 minutes), and space is in the horizontal axis (scalebar 2 μm). (**B-C**) Microtubule growth speed as a function of spindle length (B) or distance from the plus-end to the closest pole (C) at rescue (or first point if rescue could not be exactly determined). Microtubule growth events of clear duration are shown as round dots; other events are shown as stars. Thick black line represent average of binned data, error bars 95% confidence interval of the mean. (**D**) Histogram showing the distribution of the position of rescues with respect to the spindle center (dashed line). (**E**) Histogram showing the distribution of the duration of microtubule growth events. Cartoons below the axis in (B-D) illustrate how the magnitudes represented are measured. Data of 891 microtubule growth events (100 cells) from 6 independent experiments.

**Fig. 2-Supplement 1:**
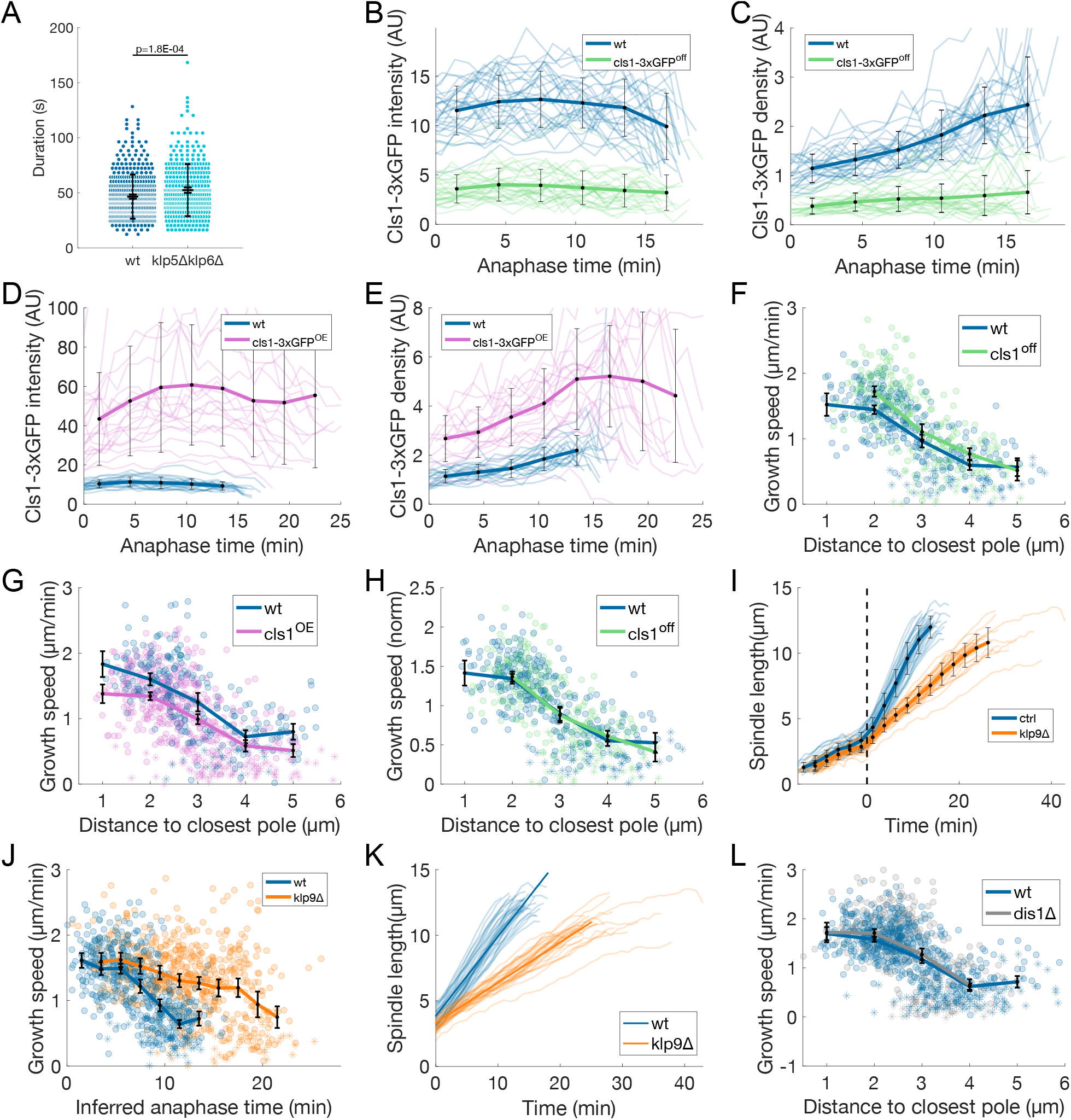
Transition from fast to slow microtubule growth occurs in the absence of known anaphase MAPs. (**A**) Microtubule growth event duration in wild-type and klp5Δklp6Δ cells. p-values correspond to T-test. Error bars represent 95% confidence interval of the mean and standard deviation. (**B,D**) Cls1-3xGFP total intensity on the spindle as a function of time after anaphase onset in wt, cls1-3xGFP^off^ (B) and cls1-3xGFP^OE^ (D) cells. (**C, E**) Total intensity of Cls1-3xGFP divided by total intensity of mCherry-Atb2 in a window of 2 μm at the spindle center as a function of time after anaphase onset in wt, cls1-3xGFP^off^ (C) and cls1-3xGFP^OE^ (E) cells. (**F-G**) Microtubule growth speed as a function of the distance from the plus-end to the closest pole at rescue (or first point if rescue could not be exactly determined) in wt, cls1^off^ (F) and cls1^OE^ (G) cells. Microtubule growth events of clear duration are shown as round dots; other events are shown as stars. (**H**) Same as (F), but microtubule growth speed is normalised to the mean growth speed in each condition. (**I**) Spindle length measured from Sid4-GFP Alp7-3xGFP signal in time (Fig. 1B), from spindle formation to spindle disassembly in wild-type and klp9Δ cells. Time is zero at anaphase onset, indicated by the dashed line. (**J**) Microtubule growth speed as a function of anaphase time at rescue (or first point if rescue could not be exactly determined), inferred from the spindle length using the polynomial fits shown in (K) (see Methods). (**K**) Anaphase spindle length as a function of time, as in (I), thin lines represent individual trajectories, thick lines represent fits to first degree polynomials of each condition used to calculate inferred anaphase time in (J) (see Methods). (**L**) Microtubule growth speed as a function of the distance from the plus-end to the closest pole at rescue (or first point if rescue could not be exactly determined) in wt and dis1Δ cells. Thin lines in (B-E, I, K) represent individual trajectories, thick lines in (B-J, L) represent average of binned data. Error bars in (B-L) represent the 95% confident interval of the mean. Number of observations: (A) 431 (47 cells) wt, 349 (29 cells) klp5Δklp6Δ microtubule growth events from 5 independent experiments (B, C) 30 wt, 31 cls1^off^ anaphase spindles from 3 independent experiments, (D, E) 21 wt, 20 cls1^OE^ anaphase spindles from 2 independent experiments, (F, H) 304 (24 cells) wt, 277 (31 cells) cls1^off^ microtubule growth events from 4 independent experiments, (G) 248 (22 cells) wt, 532 (40 cells) cls1^OE^ microtubule growth events from 3 independent experiments, (I, K) 30 wt, 30 klp9Δ spindles from 2 independent experiments, (J) 531 (36 cells) wt, 610 (40 cells) klp9Δ microtubule growth events from 4 independent experiments, (L) 614 (50 cells) wt, 367 (26 cells) dis1Δ microtubule growth events from 3 independent experiments.

**Fig. 3-Supplement 1:**
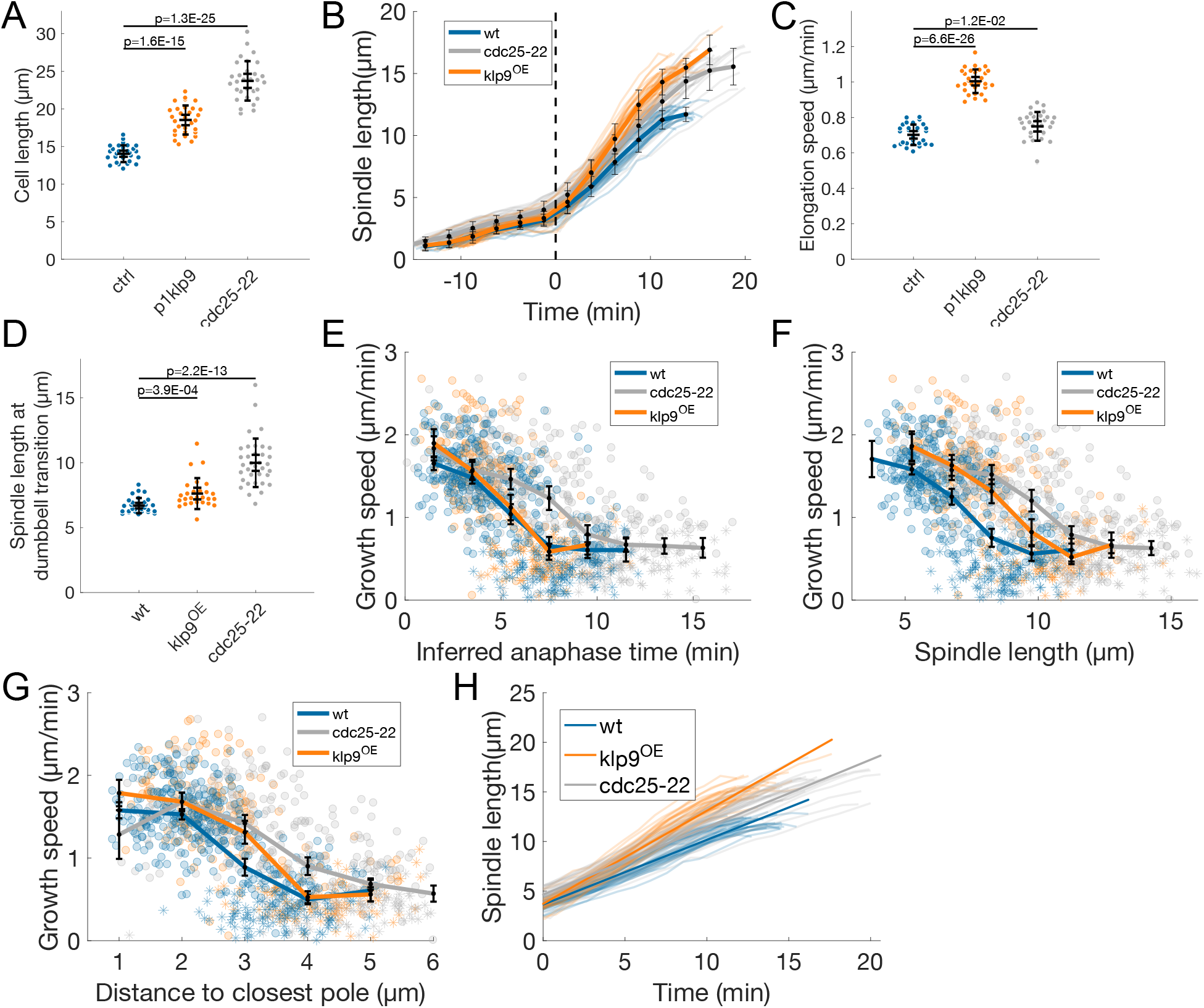
Microtubules grow slower when they enter the nuclear membrane bridge formed at the dumbbell transition. Comparison of wild-type (blue), klp9^OE^ (orange) and cdc25-22 (grey) cells. (**A**) Cell length at mitosis start. (**B**) Spindle length measured from Sid4-GFP Alp7-3xGFP signal in time (Fig. 1B), from spindle formation to spindle disassembly. Time is zero at anaphase onset (dashed line). Thin lines represent individual trajectories, thick lines represent average of binned data. Error bars represent 95% confidence interval of the mean. (**C**) Anaphase spindle elongation speed. (**D**) Spindle length at dumbbell transition. (**E, F, G**) Microtubule growth speed as a function of inferred anaphase time (E, see Methods), spindle length (F) and distance to closest pole (G) at rescue (or first point if rescue could not be exactly determined). Microtubule growth events of clear duration are shown as round dots; other events are shown as stars. Thick lines represent average of binned data, error bars 95% confidence interval of the mean. Inferred anaphase time in (E) is calculated from the polynomial fits in (H) (see Methods). (**H**) Anaphase spindle length as a function of time, as in (B), thin lines represent individual trajectories, thick lines represent fits to first degree polynomials of each condition used to calculate inferred anaphase time in (E) (see Methods). p-values in (B, C, D) correspond to T-test. Number of observations: (A-C, H) 30 cells of each condition, from one experiment. (D) 30 wt, 27 klp9^OE^, 35 cdc25-22 cells from 3 independent experiments. (E-G) 442 (30 cells) wt, 260 (27 cells) klp9^OE^, 401 (35 cells) cdc25-22 microtubule growth events from 3 independent experiments.

**Fig. 4-Supplement 1:**
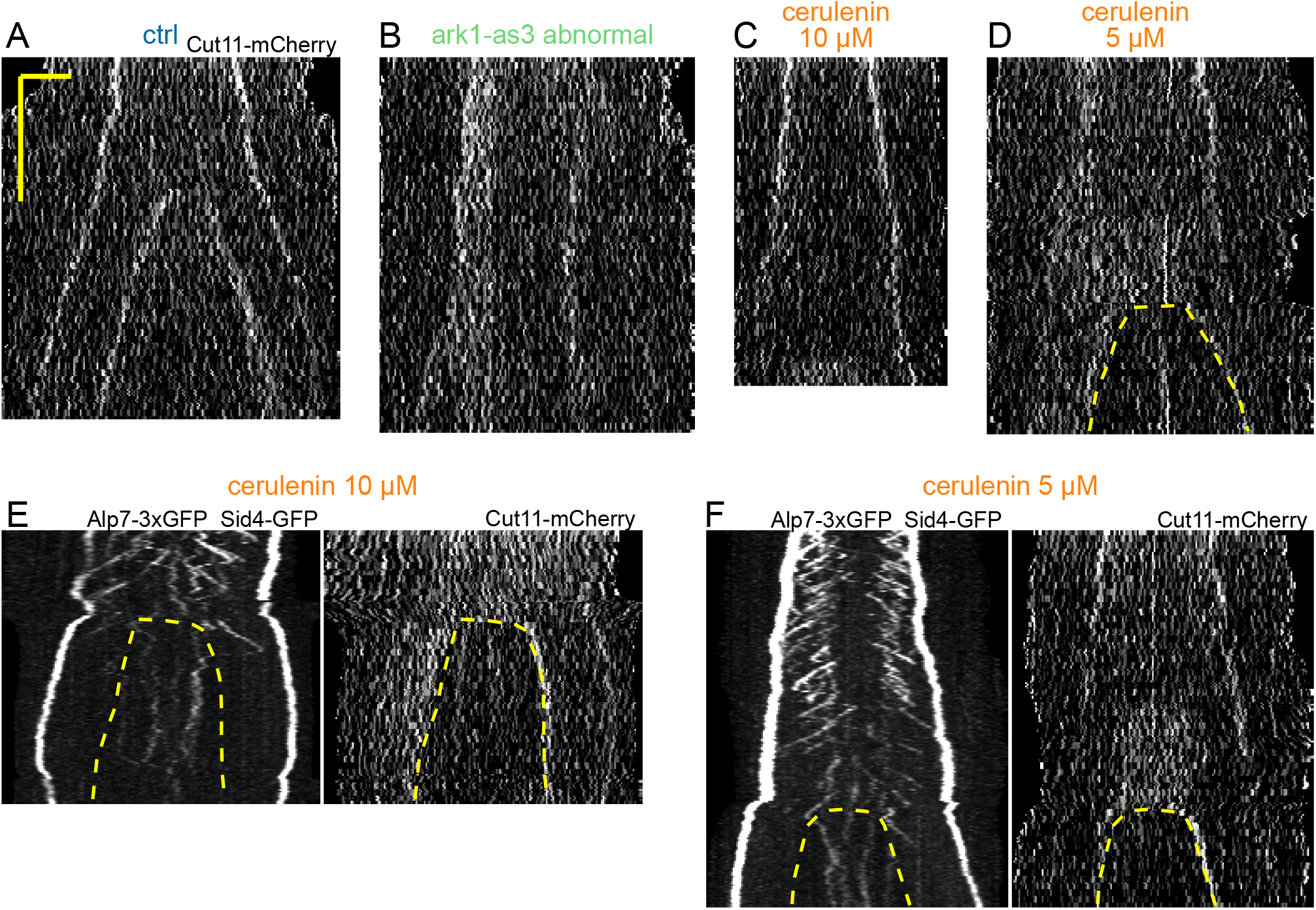
Preventing the dumbbell transition abolishes the transition from fast to slow microtubule growth. (**A-D**) Kymographs of the Cut11-mCherry channel, for the same spindles as Fig. 4E-H. Time is in the vertical axis (scalebar 5 minutes), and space is in the horizontal axis (scalebar 2 μm). (**E-F**) Kymographs of spindles that underwent dumbbell transition in the presence of 10 μM (E) and 5 μM (F) cerulenin. Scale as in A. Dashed lines outline the nuclear membrane bridge formed after the dumbbell transition.

**Fig. 5-Supplement 1:**
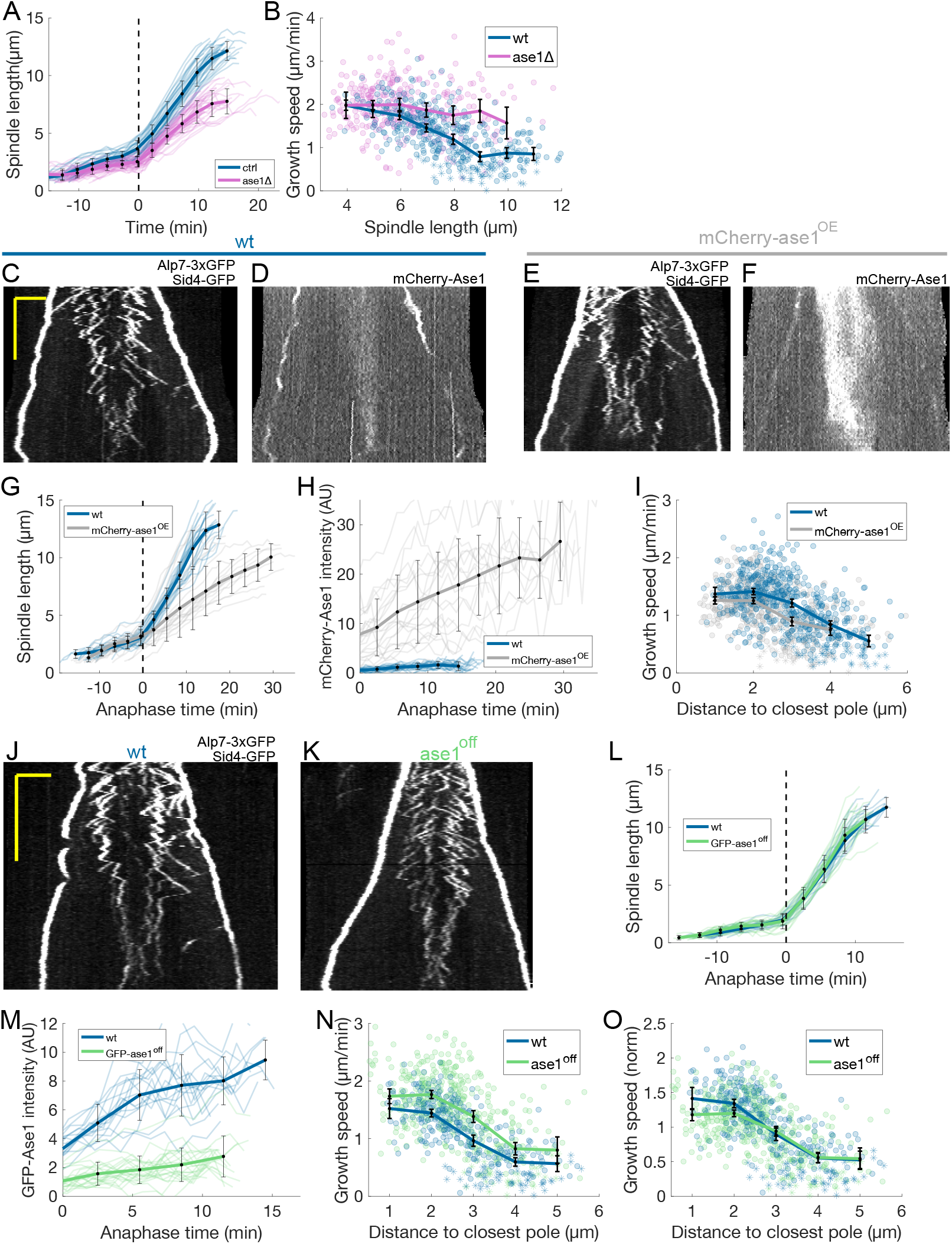
Ase1 is required for normal rescue distribution and microtubule growth speed to decrease in anaphase B. (**A**) Spindle length measured from Sid4-GFP Alp7-3xGFP signal in time, from spindle formation to spindle disassembly. Time is zero at anaphase onset. The dashed line indicates the start of anaphase. Same data as Fig. 5E. (**B**) Microtubule growth speed as a function of spindle length at rescue (or first point if rescue could not be exactly determined) in wild-type (blue) and ase1Δ (pink) cells. (**C-F**) Kymographs of anaphase mitotic spindles in cells expressing Alp7-3xGFP, Sid4-GFP and mCherry-Ase1 from its endogenous promoter (wild-type, C, D) or overexpressed under the control of a P1nmt1 promoter (mCherry-ase1^OE^, E, F). Time is in the vertical axis (scalebar 5 minutes), and space is in the horizontal axis (scalebar 2 μm). (**G**) Spindle length as a function of time in wild-type (blue) and mCherry-ase1^OE^ (grey) cells. Data represented as in (A). (**H**) mCherry-Ase1 total intensity as a function of time after anaphase onset in wild-type (blue) and mCherry-ase1^OE^ (grey) cells. (**I**) Microtubule growth speed as a function of the distance between the plus end and the closest pole at rescue (or first point if rescue could not be exactly determined) in wild-type (blue) and mCherry-ase1^OE^ (grey) cells. (**J-K**) Kymographs of anaphase mitotic spindles in in wild-type (J) and ase1^off^ (K) cells expressing Alp7-3xGFP and Sid4-GFP. Time is in the vertical axis (scalebar 5 minutes), and space is in the horizontal axis (scalebar 2 μm). (**L**) Spindle length as a function of time in wild-type (blue) and GFP-ase1^off^ (green) cells. Data represented as in (A). (**M**) GFP-Ase1 total intensity on the spindle as a function of time after anaphase onset in wild-type (blue) and GFP-ase1 ^off^ (green) cells. (**N**) Microtubule growth speed as a function of the distance between the plus end and the closest pole at rescue (or first point if rescue could not be exactly determined) in wild-type (blue) and ase1^off^ (green) cells. (**O**) Same as (N), but microtubule growth speed is normalised to the mean growth speed in each condition. Thin lines represent individual trajectories, thick lines represent average of binned data. Error bars represent the 95% confident interval of the mean. Number of observations: (A) 30 wt, 48 ase1Δ cells from 3 independent experiments (B) 402 (34 cells) wt, 316 (39 cells) ase1Δ microtubule growth events from 4 independent experiments (G, H) 20 wt, 21 mCherry-ase1^OE^cells from 1 experiment (I) 567 (50 cells) wt, 490 (43 cells) mCherry-ase1^OE^microtubule growth events from 4 independent experiments (L, M) 16 wt, 21 GFP-ase1 ^off^ cells from 2 independent experiments (N, O) 304 (24 cells) wt, 430 (41 cells) ase1^off^ microtubule growth events from 4 independent experiments

**Fig. 6-Supplement 1:**
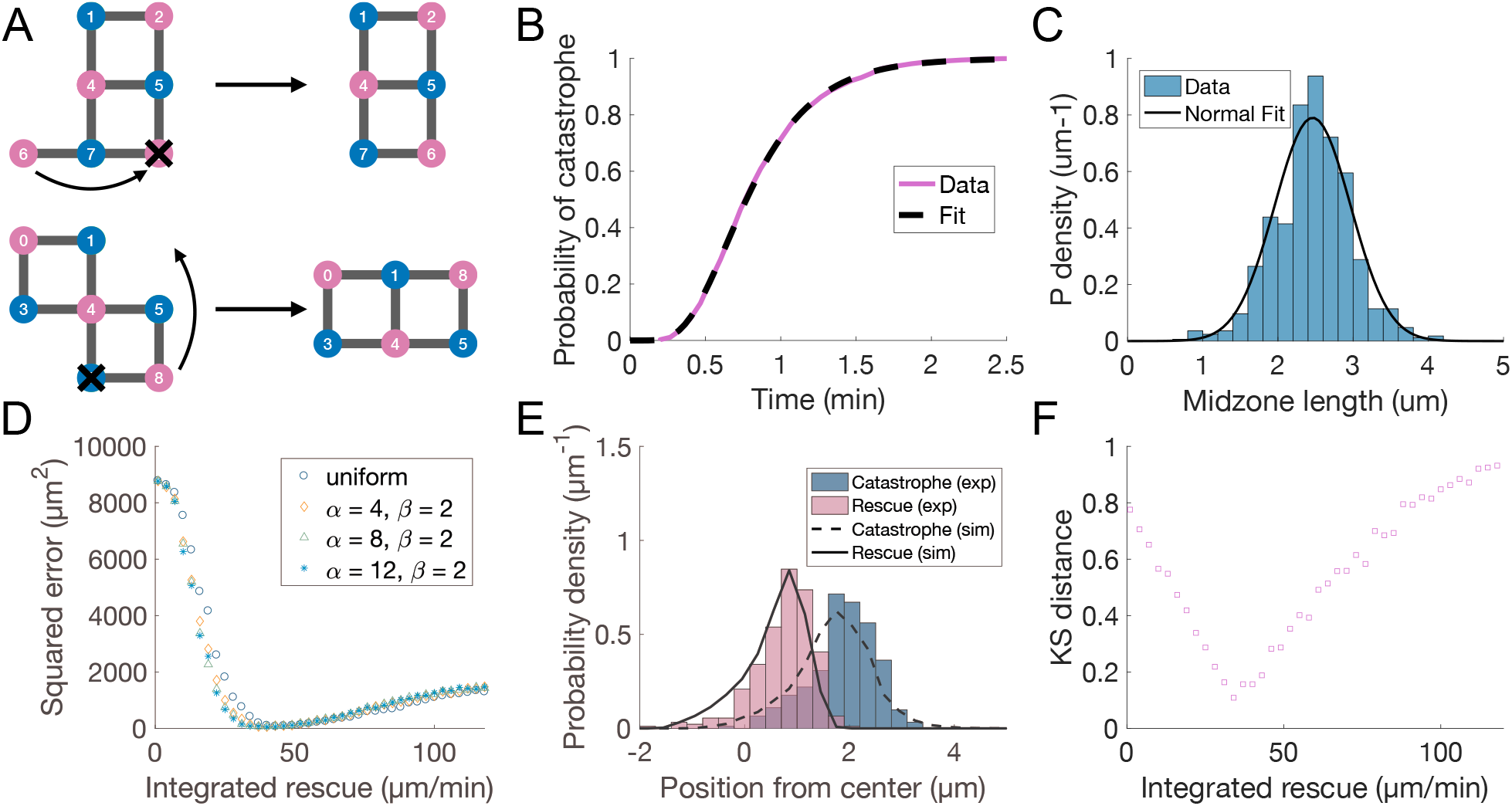
Promoting microtubule rescues at the midzone edge is sufficient to coordinate sliding and growth. (**A**) Cartoon depicting microtubule rearrangements occurring in simulations following the loss of a microtubule. They can happen in two ways. (top) A microtubule is lost and there is an existing one in the same orientation which has less neighbours. (bottom) A neighbour of a lost microtubule would have more neighbours if it was in another position. (**B**) Cumulative distribution of duration of microtubule growth events in cells expressing Sid4-GFP and Alp7-3xGFP (blue line, same data as Fig. 1-Supplement 1C), plotted along the fit of the data to Equation 4 (pink dashed line, *θ*=3.17 min^-1^ and *n*=8.53). **(C)** Histogram showing the distribution of midzone length in cells expressing mCherry-Atb2 and Cls1-3xGFP (same data as in Fig. 1G). Black line represents a fit to a normal distribution (*μ*=2.5 μm, *σ*=0.5 μm) (**D**) Sum of squared error resulting from comparing total polymer length as a function of spindle length in simulations with total tubulin intensity as a function of spindle length in experiments (pink data in Fig. 6C, see Methods). Each dot corresponds to a set of 200 simulations with equal parameters, and is placed on the x axis according to the value of integrated rescue *R*. (**E**) Distribution of positions of microtubule catastrophe and rescue with respect to the spindle center in experiments (histograms, same data as Fig. 1G), and 200 simulations (lines), for *R* = 55μm/min, *α*=1, *β*=1 (uniform rescue). (**F**) Kolmogorov-Smirnov distance obtained when comparing the distribution of spindle length at collapse in ase1Δ cells (data in Fig. 6H) and simulations. This distance can go from 0 to 1 and is smaller when the probability distributions in experiments and simulations are similar (see Methods). Each dot corresponds to a set of 200 simulations with equal parameters, and is placed on the x axis according to the value of integrated rescue *R* (Equation 5). The minimal value of Kolmogorov-Smirnov distance corresponds to *R*=34 μm/min, and this is the value used in Fig. 6G, H. See Table S2 for simulation parameters.

**Table S1:** Regression analysis and 2-way analysis of variance. Regression analysis summary and 2-way anova summary for the data represented in Fig. 3D (Top), Fig. 4I (center) and Fig. 4J (bottom). The tables are the output of the functions `fitlm` and `anova` in Matlab. The comparison is done with two categories: ‘condition’ (genetic background or drug treatment) and ‘position’ (before / outside / inside, see cartoons in Fig. 3 and 4). The reference category is microtubule growth events in the condition wild type rescued before the dumbbell transition. We also take into account the interaction between the ‘condition’ and ‘position’ variables.

|  |  |  | Estimate | SE | tStat | pValue |
| --- | --- | --- | --- | --- | --- | --- |
| (Intercept) |  |  | 1.5548 | 0.032617 | 47.669 | 2.8392e-269 |
| condition_cdc25-22 |  |  | 0.033894 | 0.053382 | 0.63495 | 0.5256 |
| condition_plklp9 |  |  | 0.10648 | 0.064349 | 1.6548 | 0.098254 |
| position_outside |  |  | -0.16789 | 0.067173 | -2.4994 | 0.012584 |
| position_inside |  |  | -0.92376 | 0.045654 | -20.234 | 1.3913e-77 |
| condition_cdc25-22:position_outside |  |  | 0.036012 | 0.10072 | 0.35756 | 0.72074 |
| condition_plklp9:position_outside |  |  | 0.068408 | 0.10451 | 0.65455 | 0.51289 |
| condition_cdc25-22:position_inside |  |  | 0.067308 | 0.068658 | 0.98033 | 0.32714 |
| condition_plklp9:position_inside |  |  | -0.032138 | 0.081503 | -0.39432 | 0.69342 |
|  | SumSq | DF | MeanSq | F | pValue |  |
| condition | 1.987 | 2 | 0.99352 | 4.9675 | 0.0071183 |  |
| position | 204.69 | 2 | 102.35 | 511.73 | 1.3234e-157 |  |
| condition:position | 0.61479 | 4 | 0.1537 | 0.76848 | 0.54578 |  |
| Error | 218.8 | 1094 | 0.2 |  |  |  |

|  |  |  | Estimate | SE | tStat | pValue |
| --- | --- | --- | --- | --- | --- | --- |
| (Intercept) |  |  | 1.4789 | 0.043995 | 33.615 | 5.4385e-163 |
| condition_ark1-as3 normal |  |  | -0.0172 | 0.075479 | -0.22788 | 0.81979 |
| condition_ark1-as3 abnormal |  |  | -0.067614 | 0.072137 | -0.93731 | 0.34884 |
| position_inside |  |  | -0.6823 | 0.053627 | -12.723 | 2.589e-34 |
| position_outside |  |  | -0.15671 | 0.058799 | -2.6652 | 0.0078279 |
| condition_ark1-as3 normal:position_inside |  |  | 0.12857 | 0.088494 | 1.4528 | 0.1466 |
| condition_ark1-as3 abnormal:position_inside |  |  | 0.59961 | 0.10436 | 5.7457 | 1.2387e-08 |
| condition_ark1-as3 normal:position_outside |  |  | -0.09415 | 0.11228 | -0.83852 | 0.40195 |
| condition_ark1-as3 abnormal:position_outside |  |  | 0.11928 | 0.090447 | 1.3187 | 0.18758 |
|  | SumSq | DF | MeanSq | F | pValue |  |
| condition | 3.0283 | 2 | 1.5142 | 7.5952 | 0.00053455 |  |
| position | 46.232 | 2 | 23.116 | 115.95 | 1.0492e-45 |  |
| condition:position | 7.9215 | 4 | 1.9804 | 9.9336 | 7.1201e-08 |  |
| Error | 186.2 | 934 | 0.19936 |  |  |  |

|  |  |  | Estimate | SE | tStat | pValue |
| --- | --- | --- | --- | --- | --- | --- |
| (Intercept) |  |  | 1.5271 | 0.030487 | 50.092 | 4.9906e-232 |
| condition_cerulenin |  |  | 0.13681 | 0.038233 | 3.5783 | 0.00037003 |
| position_inside |  |  | -0.89616 | 0.047154 | -19.005 | 4.319e-65 |
| position_outside |  |  | -0.12986 | 0.062296 | -2.0845 | 0.037482 |
| condition_cerulenin:position_inside |  |  | -0.41416 | 0.13992 | -2.9599 | 0.0031826 |
| condition_cerulenin:position_outside |  |  | -0.39908 | 0.29753 | -1.3413 | 0.18027 |
|  | SumSq | DF | MeanSq | F | pValue |  |
| condition | 1.2695 | 1 | 1.2695 | 7.5462 | 0.0061704 |  |
| position | 79.034 | 2 | 39.517 | 234.9 | 1.5484e-78 |  |
| condition:position | 1.7317 | 2 | 0.86587 | 5.1471 | 0.0060418 |  |
| Error | 116.08 | 690 | 0.16823 |  |  |  |

**Table S2:** Simulation parameters.

| Symbol | Meaning | Value | Source |
| --- | --- | --- | --- |
| $v_p$ | Microtubule growth speed | 1.6 $\mu\text{m}/\text{min}$ when fixed, uniformly sampled between 0.35 and 1.5 $\mu\text{m}/\text{min}$ when scanned | Fig. 1F |
| $v_d$ | Microtubule shrinking speed | 3.6 $\mu\text{m}/\text{min}$ | Fig. 1-Supplement 1D |
| $v_s$ | Microtubule sliding speed | 0.35 $\mu\text{m}/\text{min}$ | Fig. 1-Supplement 1A |
| $R$ | Integrated rescue rate | 55 $\mu\text{m}/\text{min}$ when fixed for wild type, 34 $\mu\text{m}/\text{min}$ when fixed for $\text{ase1}\Delta$, uniformly sampled between 1 and 120 $\mu\text{m}/\text{min}$ when scanned | |
| $n$ | See Equation 4 | 8.53 in wild type, 6.8 in $\text{ase1}\Delta$ | Fig. 6-Supplement 1B for wt, data not shown for $\text{ase1}\Delta$ |
| $\theta$ | See Equation 4 | 3.17 $\text{min}^{-1}$ in wild type, 2.5 $\text{min}^{-1}$ in $\text{ase1}\Delta$ | Fig. 6-Supplement 1B for wt, data not shown for $\text{ase1}\Delta$ |
| $\mu$ | Mean of normal fit | 1.23 $\mu\text{m}$ | Fig. 6-Supplement 1C |
| $\sigma$ | Standard deviation of normal fit | 0.25 $\mu\text{m}$ | Fig. 6-Supplement 1C |
| $\alpha$ | Parameter of beta distribution | 1 in uniform, indicated in legend otherwise | |
| $\beta$ | Parameter of beta distribution | 1 in uniform, 2 otherwise | |
| $h$ | Simulation timestep | 0.01 min | |

**Movie 1:**
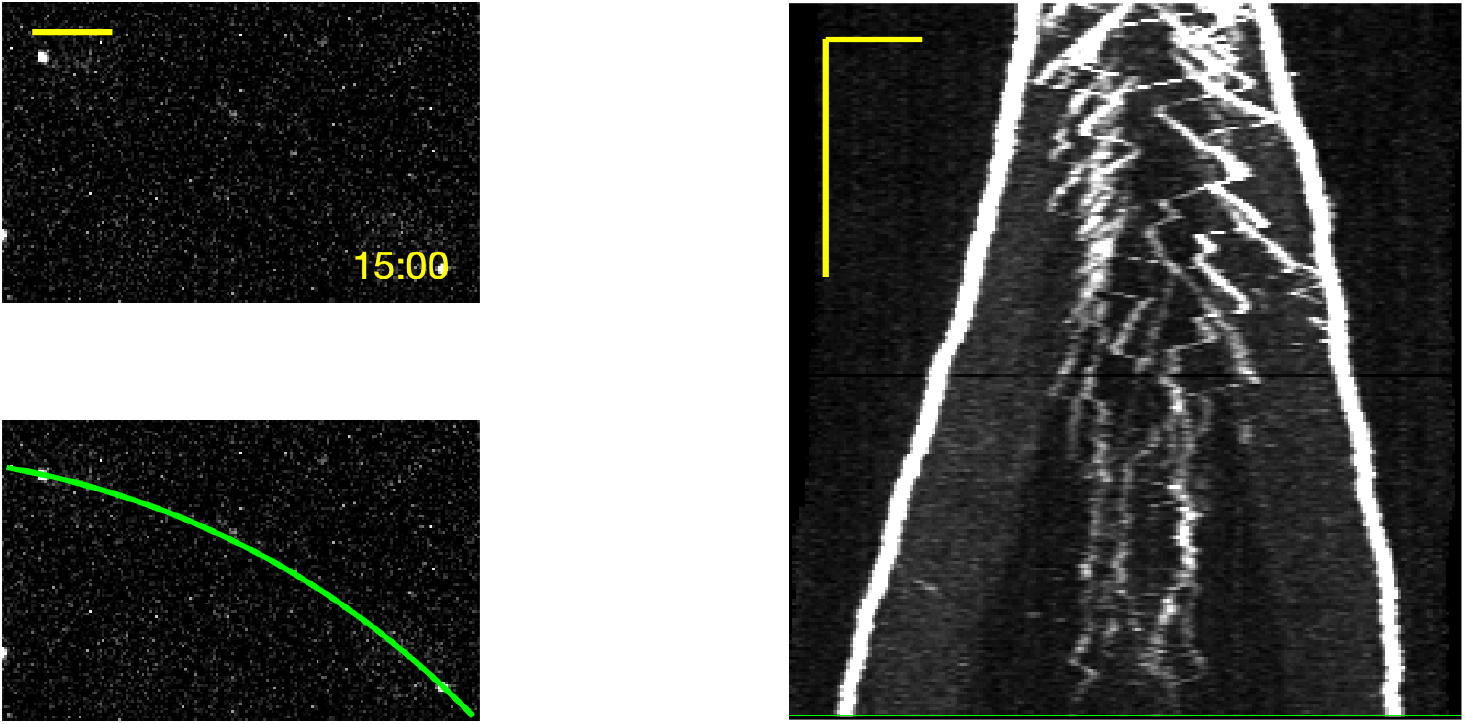
Kymograph construction in cells expressing Alp7-3xGFP Sid4-GFP. Construction of the kymograph shown in Fig. 1C from a live-imaging movie of cells expressing Sid4-GFP and Alp7-3xGFP The green curve in the movie marks the fitted spindle trace (a second order polynomial) used to obtain an intensity profile and produce the kymograph shown on the right. In the kymograph, time is in the vertical axis (scalebar 5 minutes), and space is in the horizontal axis (scalebar 2 μm). The time on the top left movie is in minutes:seconds, scalebar in the movie is 2 μm.

